# A fundamental law underlying predictive remapping

**DOI:** 10.1101/2023.01.24.525276

**Authors:** Ifedayo-EmmanuEL Adeyefa-Olasupo

## Abstract

Predictive remapping (*R*) — the ability of cells in retinotopic brain structures to transiently exhibit spatiotemporal shifts beyond the spatial extent of their classical anatomical receptive fields — has been proposed as a primary mechanism that stabilizes an organism’s percept of the visual world around the time of a saccadic eye movement. Despite the well-documented effects of *R*, a biologically plausible mathematical abstraction that specifies a fundamental law and the functional architecture that actively mediates this ubiquitous phenomenon does not exist. I introduce the Newtonian model of *R*, where each modular component of *R* manifests as three temporally overlapping forces - a centripetal 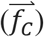, convergent 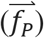 and translational force 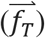, that perturb retinotopic cells from their equilibrium extent. The resultant and transient influences of these forces 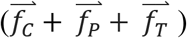 gives rise to a neuronal force field that governs the spatiotemporal dynamics of *R*. This neuronal force field fundamentally obeys an inverse-distance law, akin to Newton’s law of universal gravitation [1] and activates retinotopic elastic fields (elφs). I posit that elφs are transient functional structures that are self-generated by a visual system during active vision and approximate the sloppiness (or degrees of spatial freedom) within which receptive fields are allowed to shift while ensuring that retinotopic organization does not collapse. The predictions of the proposed general model are borne out by the spatiotemporal changes in sensitivity to probe stimuli in human subjects around the time of a saccadic eye movement and qualitatively match neural signatures associated with predictive shifts in the receptive fields of cells in premotor and higher-order retinotopic brain structures.

## I. INTRODUCTION

In biological systems that possess a fovea, the central 2° of visual space is extensively represented in the retina, premotor, and primary visual brain structures, and is therefore well suited for the detection and detailed inspection of ecologically relevant visual stimuli. During active vision, directed saccadic eye movements are constantly recruited by foveated visual systems to bring new stimuli of interest that fall on the peripheral retina into the central 2° of visual space [2–3]. In foveated systems and afoveated systems (i.e., a visual system that represents visual space in a uniformed manner) directed saccadic eye movements cause large and rapid displacements of the retinal image. These displacements introduce significant retinal disruptions, akin to those observed when attempting to photograph a rapidly moving stimulus using a camera. Indeed, different visual systems have evolved to account for these sudden retinal image disruptions such that their gaze is quickly stabilized, and their perception of the visual world remains relatively continuous [4–7]. The ability of a visual system in general to account for these incessant retinal disruptions is commonly referred to as perceptual (or spatial) constancy [8–12].

Translational remapping — the ability of cells within retinotopic brain structures to predictively shift their neural sensitivity beyond the spatial extent of their classical anatomical receptive fields to their future post-saccadic locations just before saccade onset — has been proposed as a primary mechanism that mediates spatial constancy (Supp. Fig) **1A** [13–16]. However, more recent electrophysiological studies have challenged this dominant view of predictive remapping (R) and have instead demonstrated that these transient predictive shifts are directed towards and around a preselected peripheral site which includes the location of the saccade target (Supp. Fig) **1B**. This form of R is referred to as convergent remapping [17–19].

Translational and convergent accounts of R thus remain at odds, and the functional role that these divergent forms of transient receptive field shifts play in mediating spatial constancy is vigorously debated [20–23]. For example, prior studies have routinely sampled regions in space that yield results favorable to one form of R over another. In fact, with unbiased spatial sampling, a more recent study suggests that the functional correlates of convergent remapping may include additional components [24], an account that contradicts earlier results regarding the functional correlates of R [25–26]. In addition to this, the only study that did increase spatial sampling around a preselected peripheral site was devoid of a computational framework to correctly contextualize and interpret reported observations [27]. It is therefore not surprising that the observations from this study are radically inconsistent with well-established classical neurophysiological accounts of R [13–16].

Equally unresolved is the issue of time. First, most studies have only assessed the consequences of R on visual sensitivity within a limited pre-saccadic window (e.g. −250ms to +50ms with respect to the onset of a saccadic eye movement). To this end, it is unclear how transient receptive field shifts that occur within earlier pre-saccadic windows actively modulate peri-saccadic and post-saccadic sensitivity [23]. Finally, a recent study has reported that R includes a translational component followed by a convergent component, with each component specified within distinct non-overlapping temporal windows [28–29]. This temporal account of R is however at odds with the canonical order of pre-saccadic events [8] and a more recent study has challenged these results on methodological grounds [19].

In addition to these unresolved issues, there are three important questions about the visuomotor system previous investigations have overlooked. First, the frequency of saccades (~2-4 times per second) [30] and the accompanying predictive receptive field shifts can be energetically expensive [31]. Yet, a fundamental law visual systems in general use to balance energy costs with predictive function remains a mystery. Second, the functional architecture that supports increased or sustained sensitivity of retinotopic cells beyond their anatomical classic center-surround structure, while preventing radical and unsustainable forms of remapping (e.g., over-convergence or over-translation) is unknown. Finally, no computational investigations of R have been able to provide a mechanistic underpinning that ensures the immediate availability of neural resources at the future post-saccadic location of the saccade target [32]. This rapid availability of neural resources cannot be explained by pure translational or convergent shifts.

To provide an account for (a) the spatiotemporal characteristics that define R, (b) a fundamental law and the functional architecture which ensures that retinotopic cells are appropriately sensitized towards different locus in visual space while preserving retinotopic organization around the time of a saccadic eye movement, and (c) the jump in late pre-saccadic sensitivity in the periphery and the immediate availability of neural resources at the future center of gaze, I developed a simple yet powerful general model of R derived using Newtonian mechanics. Next, I simulated population neural density signatures. I assume that these density signatures are equivalent to visual and neural sensitivity readouts that have been reported in previous functional and neurophysiological studies [18, 19]. I later used these signatures to assess the extent they could either predict or qualitatively match previous functional and neurophysiological studies. Finally, to directly assess the generalizability and biological plausibility of the model, I conducted three psychophysics experiments. These experiments allowed me to directly investigate the transient consequences of R on visual sensitivity to flashed visual probe stimuli in human subjects within foveal, parafoveal, and peripheral locations in visual space, and a larger pre- and post-saccadic window (−~600ms to + 150ms), results which I later compared to stimulated results. Comparisons between experimental and simulated results was followed by rhythmic investigations used to further scrutinize a core assumption of the model.

## II. MODEL

Since the initial discovery of R ~45 years ago [33], an assortment of models has attempted to specify a biophysical mechanism, a specific retinotopic brain structure, and or a neural circuit underlying this ubiquitous phenomenon [34–38]. Models that opted for a less reductionist approach were carefully parametrized and or included non-modular components, with simulations ultimately conforming to an experimental demand (i.e., one form of remapping over another, or the combination of both) [39–42]. Indeed, these approaches, as recent reviews demonstrate [20–23] have fundamentally obscured the emergent simplicity and elegance of the phenomenon [43].

Acutely aware of the pitfalls previous models have been unable to avoid, the proposed model at present is a mathematical abstraction. By definition, such abstraction is agnostic to a specific biophysical mechanism, a specific retinotopic brain structure, or a specific neural circuitry. The proposed model is also not beholden to specific experimental demands or motivated by empirical observations from a single type of biological system (i.e., foveated systems).

The principal functional unit of the proposed model is retinotopic cells. These cells subserve retinotopic brain structures and receive either direct or indirect inputs from the retina. I will refer to the excitatory extent of these units as retinotopic receptive fields - *RF_i_s* (Fig.) **1.**

**Figure 1.**
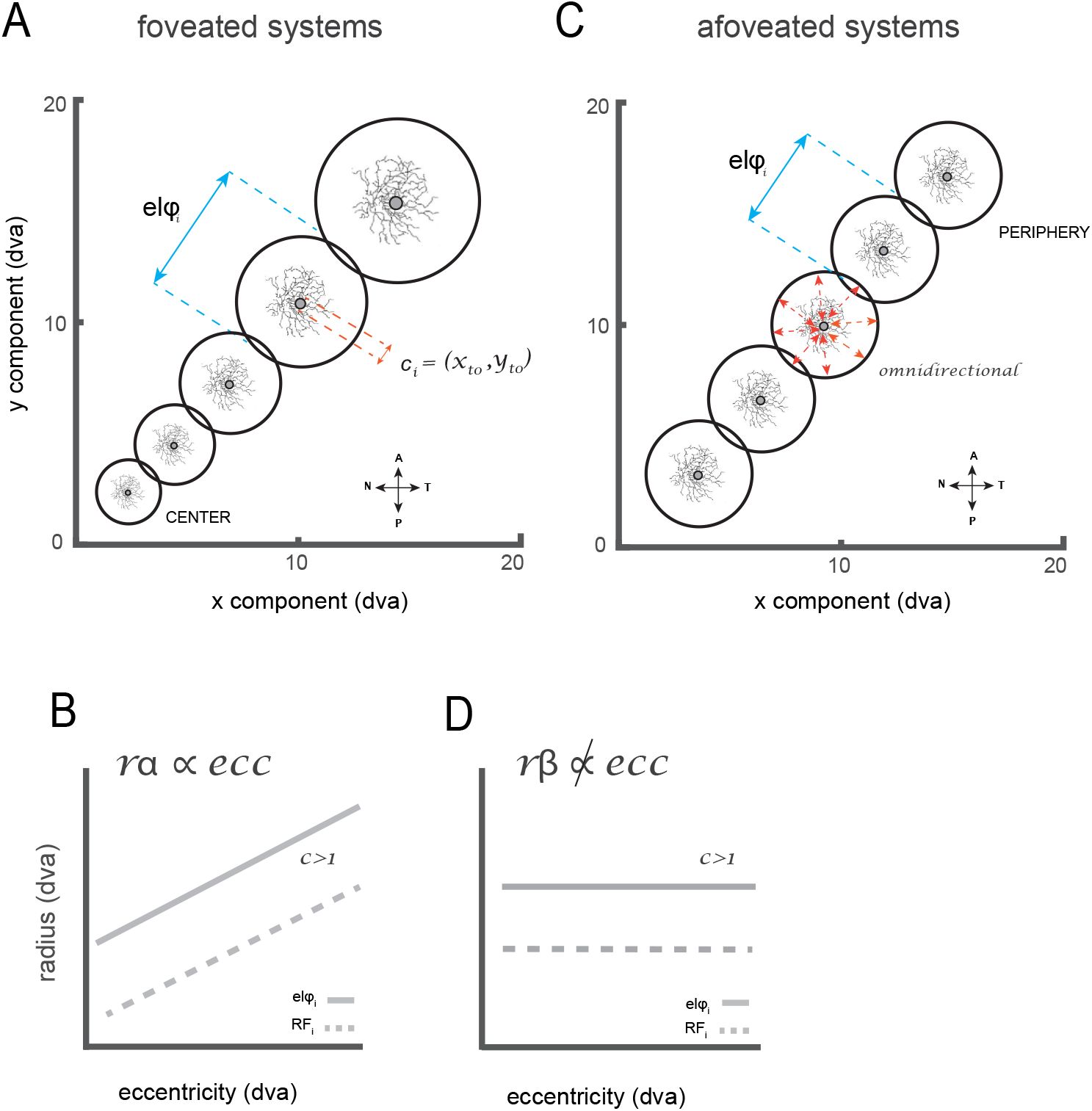
Proposed putative functional architecture that supports active vision. For every receptive field (RF_i_) there is an elastic field (elφ_i_, black annulus). Here titling of overlapping elφs along a 45 deg extent in a visual field is shown for illustration purposes. Under normal conditions RF_i_s and elφs at t_0_+1 tile the entire visual field. Depending on the biological system RF_i_s can either be: (A) eccentricity dependent, (C) non-eccentricity dependent. N, Nasal; A, anterior; T, temporal; P, posterior. (B & D) The radius of RF_i_s (in degrees of visual angle) as a function of their distance from the center of visual space (or eccentricity). When RF_i_s are multiplied by c >1, elφs will demonstrate a larger radius.

Across different retinotopic brain structures and different types of biological systems (i.e., foveated and afoveated systems), *RF_i_s* support the detection and the close inspection of ecologically relevant visual stimuli subtended within the central, intermediate, and or peripheral regions of visual space. An *RF*_i_ is characterized by two principal parameters: (a) *c_i_*, the most central location of the *RF_i_* (*x_o_*, *y_o_*) at *t_0_* (when a visual system is at an equilibrium state) (Fig) **1A**, **1C**. *c_i_* determines the *RF_i_*’s eccentricity (*ecc_i_*), that is, an *RF_i_*’s spatial extent with respect to the center of visual space, and (b) *rα*_i_, the *RF_i_*’s radius in a foveated system, which is dependent on *ecc_i_* and *k* (Eq) **2**, or *rβ_i_*, an *RF_i_*’s radius in an afoveated system which is independent of *ecc_i_* (Eq) **2**.

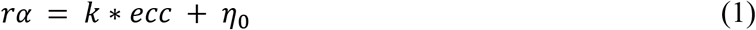

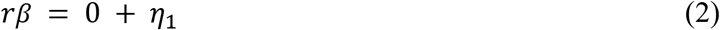

Traditionally, an *RF_i_* is thought to be constrained by its anatomical equivalent, with its *x_i_*, *y_i_* coordinate components equally fixed in retinotopic space. [44–45]. Here however I posit that anatomical classical receptive fields and their functional counterpart - *RF_i_s*, do not share a one-to-one mapping [46], specifically around the time an organism is about to reorient itself within its environment using either: a saccadic eye, head, or body movement. To this end, I further posit that *RF_i_*s fundamentally behave like spring-loaded sensors [47] that can undergo transient spatiotemporal shifts, consistent with the translational and convergent forms of R [13–19, 28, 29], and *RF_i_* dynamics that have been observed across foveated and afoveated systems [46–52]. The proposed model assumes that there are limits to which *RF_i_*s can transiently respond (or shift) beyond their classical anatomical extent such retinotopic organization is maintained at all times [47]. I, therefore, extend the idea of a retinotopic cell’s anatomical receptive field by introducing the concept of its elastic field (elφ) (Fig) **1B**, **1D.** Conserved across foveated and afoveated systems is the principle that ρ, the radius of a given elφ is the product of *rαβ* multiplied by *c*, a constant factor greater than 1 (Eq) **3**–**5.**

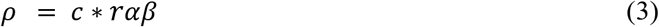

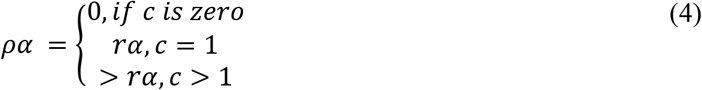

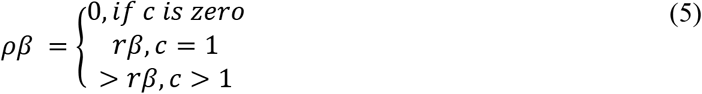

Within this mathematical range, elφs span the region immediately beyond the cell’s *c_i_*, and they are omnidirectional with respect to their classical anatomical extent (Fig)**1A, D**. elφs are transiently generated by a visual system around the time of an impending saccade, for example, a saccadic eye movement. *ρ* is fixed in time and constitute the spatial limits which *RF_i_*s are allowed to shift within. These transient spatiotemporal shifts can indeed be localized to a specific region within an elφ [46], which will cause the *RF_i_* to appear as if it has shrunk in that moment in time [17–19, 49]. Shifts well beyond the cell’s classical anatomical extent can also occur which is consistent with the translational account of R [13–16, 50]. Taken together, these transient spatiotemporal shifts manifest as distortions of population density readouts across *RF_i_*s [18,19] and temporarily distort the representation of visual space [55–59].

The onset of a retinal image evokes visually transient signals within the retina. These signals are sent to relevant retinotopic brain structures during a detailed inspection of the current stimulus, the selection of a future stimulus, and around the time of a directed or spontaneous saccadic eye movement [60] (Fig) **2A**. I use the term “active” to denote the frequency at which these signals are being sent to relevant retinotopic brain structures. The inspection and or selection of a visual stimulus before the execution of a directed or spontaneous saccadic eye movement is marked by an array of visually mediated behavioral dynamics (e.g., small fixational eye movements [61, 62], tilting of the head to visually inspect a stimulus [63, 64] These dynamics are supported by delta rhythmicity (2 to 4Hz), which is known to predict attentional shifts before a saccadic eye movement in foveated systems [65]. In response to the onset of a visual stimulus or the selection of a new one, in both foveated [33, 65–69] and afoveated systems [68], a quick burst in neural activity within pre-motor brain structures occur. I posit that these quick burst in neural activity are equivalent to oculomotor command signals and is made available to retinotopic brain structures around the time of a saccadic eye movement. I refer to the availability of these signals as “retinal image displacement signals” (Fig) **2A**. Retinotopic brain structures go on to use information about the impending displacement of the retinal image to modify persistent incoming visually transient signals originating from the retina. I refer to the modification of these signals as “feedback transient signals”. Because of the omnipresent nature of these retinal image displacement signals [70], I hypothesize these signals along with early and later feedback transient signals (both supported by delta rhythmicity) [65] uniquely equip an *RF_i_* with the ability to perform basic trigonometric operations, that is, estimate from its perspective the direction and magnitude it should shift beyond its classical anatomical extent [71]. I will refer to these early and late feedback transient signals and retinal image displacement signals, in aggregate, as “attentional-oculomotor” signals.

**Figure 2.**
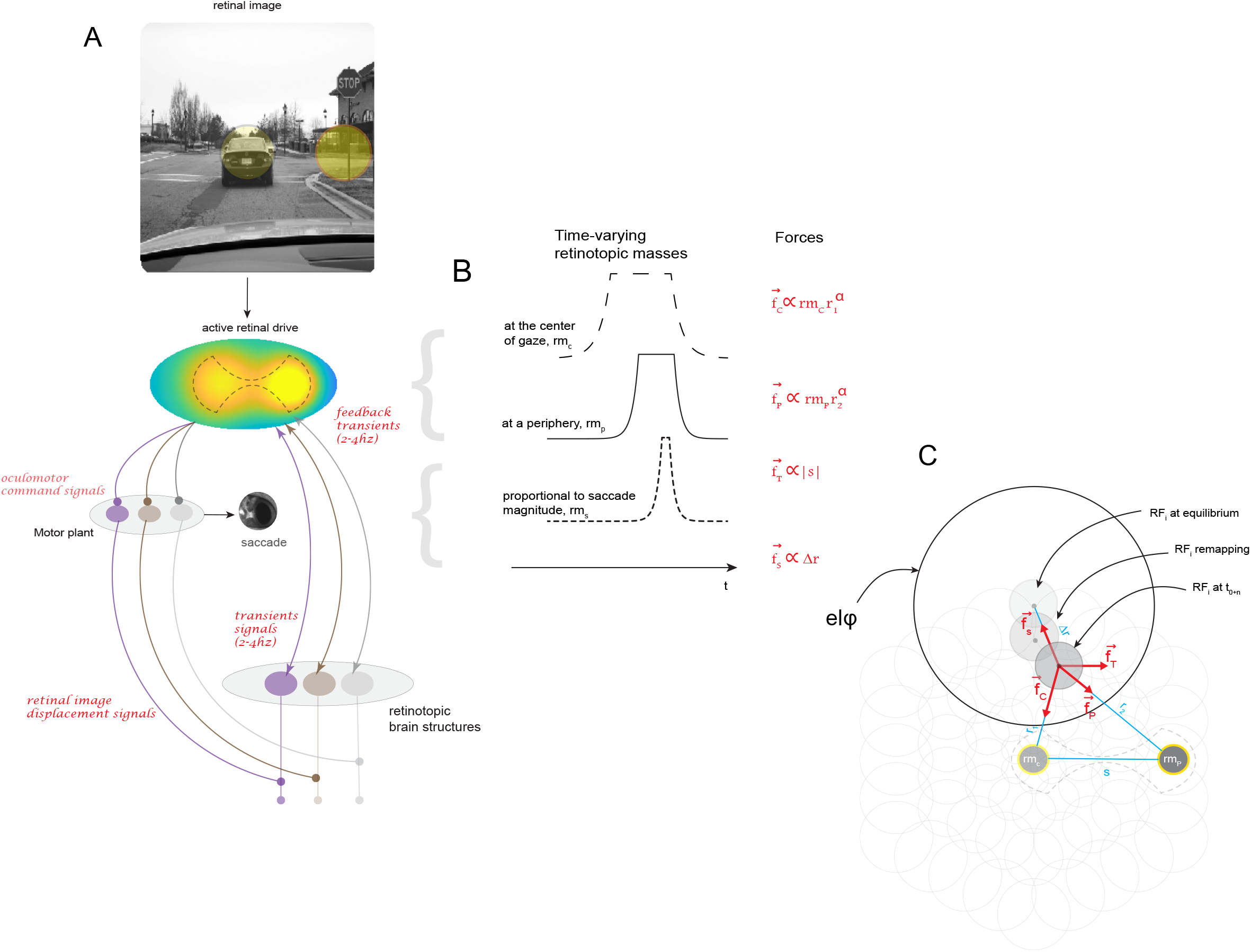
The physics underlying predictive remapping. (A) The dynamics that underlie attentional-oculomotor” signals across retinotopic brain areas is entrained by delta rhythmicity (2 to 4Hz) (B) Attentional and oculomotor signals manifest as three principal retinotopic masses. rm_C_ represents centripetal signals and is modeled as a varying mass located at the center of gaze. rm_P_ represents convergent signals and is modeled as a varying retinotopic mass located at the peripheral site corresponding to the saccade target. rm_T_ represents translational signals and is modeled as a virtual retinotopic mass at infinity located in the direction of the impending retinal displacement the saccade will induce. (C) Retinotopic field consisting of population receptive field (RF_i_s) that tile foveal, parafoveal, and peripheral regions of visual space perturbed by a force field. A single RF_i_ and its remapping extent is exaggerated in gray for illustration purposes.

At its core, the proposed model assumes that “attentional-oculomotor signals” act as forces that perturb spring-loaded *RF_i_s* from their equilibrium positions. I abstract these neurobiologically-inspired forces as being exerted by three principal time-varying retinotopic masses *rm*(*t*), subtended either at: a central (*rm_c_*) or peripheral (*rm_p_*) site in visual space, or at infinity (*rm_s_*) whose action is to produce a force in the direction of and parallel to the impending retinal displacement the saccadic eye movement will introduce. I assume that the current (*rm_c_*) or the future (*rm_p_*) location in visual space that is sampled on the retina, does not abruptly appear, and then disappear akin to a step function, but gradually appears and fades [72, 8–12]. For this very reason, I modeled each *rm_i_* as a piecewise function Eq (**6**–**8**), where *rm_i_* approaches 0 as *t* remains less than *t*_*i*1_, followed by a gradual growth (*ct^i^*) which is dependent on *i*, a stable plateau, followed by a gradual decay (*ct*^−*i*^), as *i* remains greater than 0.

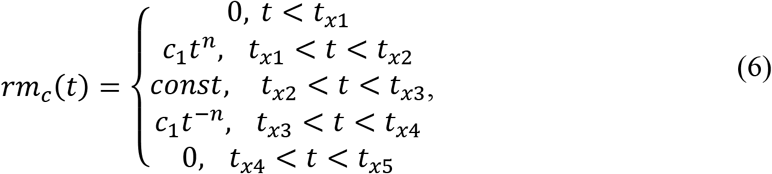

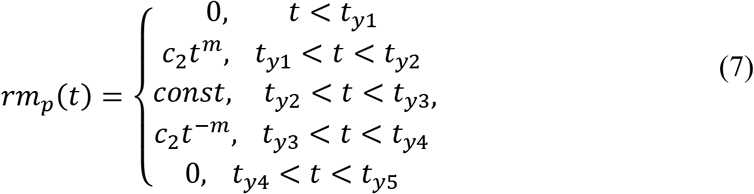

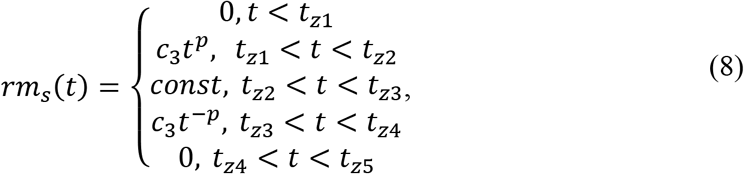

To perturb an *RF_i_*, let 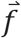 be the force that exerts its influence on an *RFi* which is the product of *rm* and 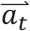, the acceleration at time *t*, where *rm* assumes unit retinotopic mass of *RF_i_* (Eq) **9**. 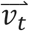 denotes the velocity at time *t* which is used to compute 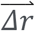, the remapping vector for an *RF_i_* from its current position (Eq) **10**–**11**. 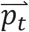 denotes the remapping operator which causes an *RF_i_* to displace its extent while being modulated by its elφ, while *r*_0_ denotes the Euclidian distance of an *RF_i_’s* current location from its anatomical extent (Eq) **12**–**13**. Taken together, the movement of an *RF_i_* from and back towards its anatomical extent produces retinotopic spatiotemporal trajectory fields Fig (**3**).

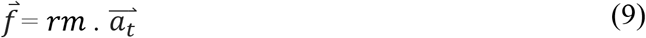

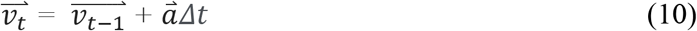

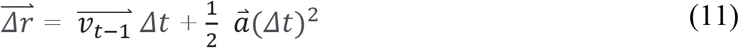

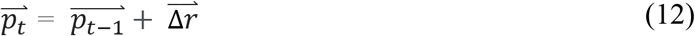

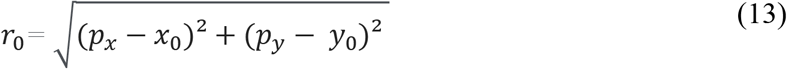

**Figure 3.**
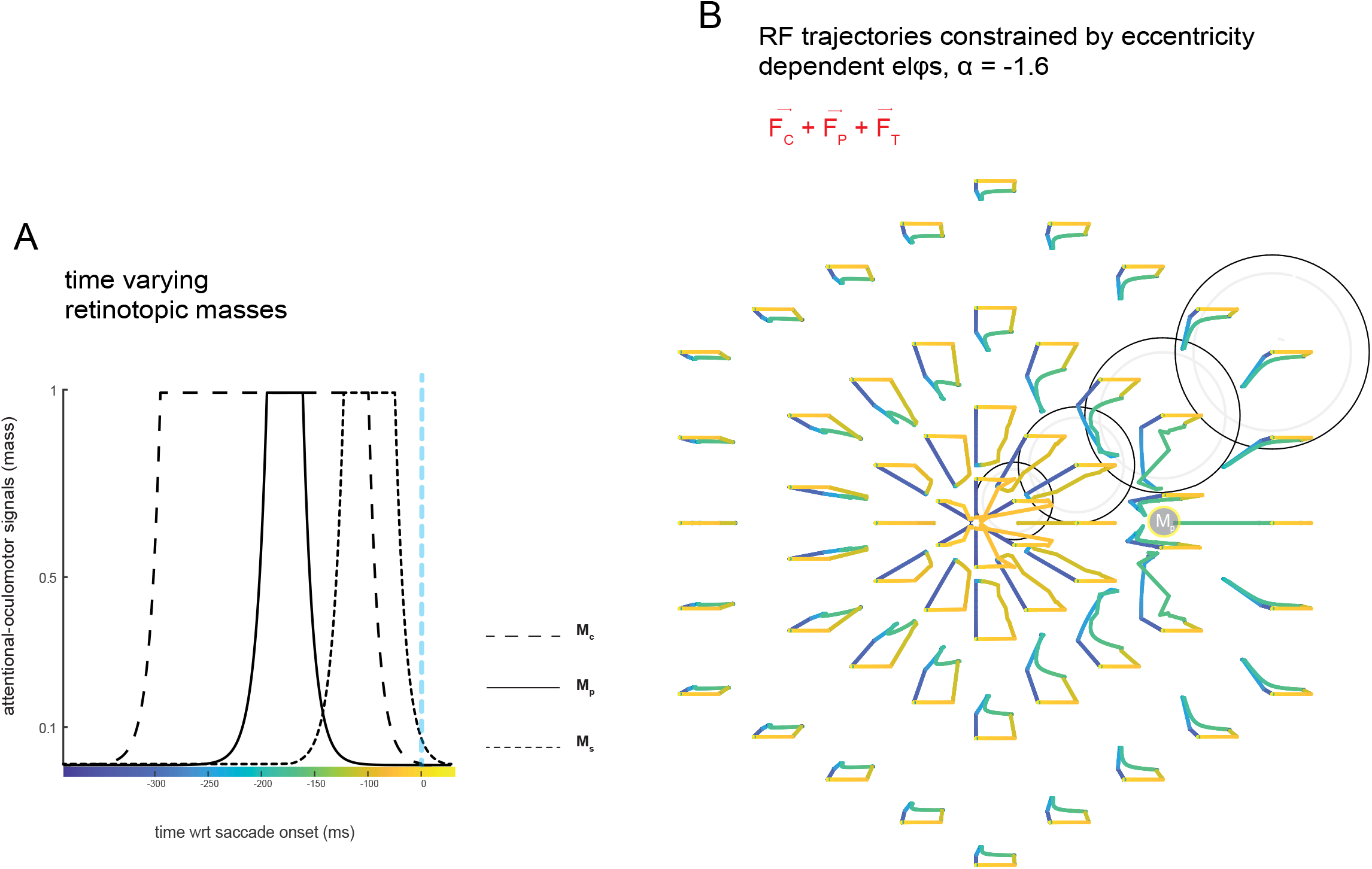
RF_i_ shifts under an inverse force field driven by simulated centripetal, convergent and translational signals. (A) Corresponding time varying masses for the case when RF_i_s were perturbed the linear combination of three types of retinotopic signals modelled as independent forces. The blue dotted line indicates the offset of the centripetal force, which marks the onset of a saccadic eye-movement. (B) spatiotemporal RF_i_ trajectories starting from the most central point of each RF_i_ under an inverse force field (α = −1.6). Note that each RF_i_ (light gray) possess an eccentricity dependent elφ, however for illustration purposes only a few are shown (black annulus). The trajectories are color coded as in (A)

I posit that pre-saccadic attentional and oculomotor processes manifest in three specific types of forces that temporally align with: (a) early visually transient signals originating from the retina, (b) late feedback trainsets that actuate the selection of a peripheral site, and (c) retinal image displacement signals derived from oculomotor motor command signals and impinge on retinotopic brain structures. The proposed model assumes that the transient perturbations of *RF_i_s*, triggered by the linear summation of these forces, is the underlying causes of R, Eq (**19**). The earliest force – a centripetal one 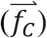 overlapping with the time of early visually transient signals during fixation – causes *RF_i_*s to transiently exhibit centripetal shifts towards the current center of gaze, where *kc_α_* is the centripetal constant, 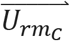 is the unit vector in the direction of *r*_1_, the spatial difference between *RF_i_s* and *rm_c_* raised to a scalar α – distance (or predictive) exponent (Eq) **14**. A centripetal force allocates a disproportionate amount of a visual system’s neural resources toward the current center of gaze [73]. This initial force is followed by a convergent force 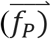, where *kp_α_* is the convergent constant, 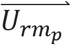 is the unit vector in the direction of *r*_2_, the spatial difference between *RF_i_s* and *rm_p_* raised to a scalar α (Eq) **15**. The onset of a convergent force precedes impending retinal image displacement signals and causes *RF_i_*s to exhibit convergent shifts towards the periphery. Counter to the dominant intuition in the extant literature [25], the principal aim of these convergent shifts is not to directly provide peripheral locations with a neural advantage when compared to less peripheral extents but divest intermediate parafoveal *RF_i_*s of their remaining resources they did not initially allocate towards the current center of gaze.

As the organism prepares and plans to execute an impending saccadic eye movement, oculomotor command signals which trigger retinal image displacement signals give rise to a translational force 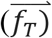, where *kt_α_* is the translational constant, 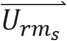 is the unit vector in the direction of *rm_s_*, while |s| is the magnitude of the impending saccade (Eq) **16**. Preceding convergent force in combination with the onset of a translational force causes *RF_i_s* to exhibit convergent and translational shifts towards a preselected peripheral site (i.e., jump-start the allocation of neural resources towards future post-saccadic locations) [18]. The growing influence of a translational force and a declining convergent force causes *RF_i_s* to directly jump towards their future post-saccadic locations [15], while a declining centripetal force (i.e., the onset of transient feedbacks) triggers convergent *RF_i_* shifts around a pre-selected target [19]. Finally, It is worth noting that the spring force, 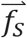, is the product of *r_o_* and the *k_0_*, the spring constant (Eq) **17**. While an elφ ensures that *RF_i_s* approaching their elastic limits are inhibited, 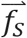 actuates the offset of each principal force. To obtain *k_o_* and *kt_α_*, the average of 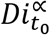 across *RF_i_*s is obtained for a given *α* with its reciprocal later computed (Eq) **18**. This produces a known range of the magnitude for 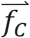 and 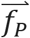 and with this *k_o_* and *kt_α_* is selected to ensure 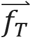 and 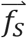 are in the same order of magnitude as 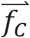 and 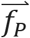.

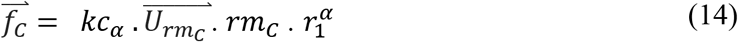

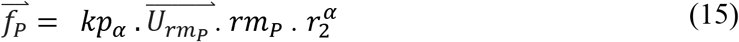

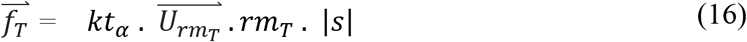

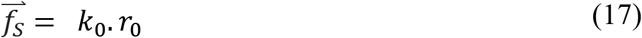

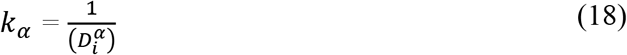

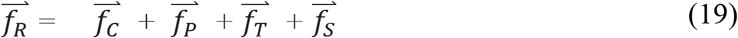

## III. RESULTS

### A. SIMULATIONS

All of the previous functional and neurophysiological studies that have explored the spatiotemporal dynamics that could characterize R were conducted using foveated biological systems. This has two very important consequences. One, the simulation results I will use here to systematically assess the phenomenological validity of the proposed model are limited to the extant literature. Two, as all rigorous models should strive towards, reported simulations will provide a rich and clear set of novel predictions that should inform future investigations of R, especially those that are conducted using afoveated systems.

In the subsequent sections, despite proposing the underlying causes of R, modular components included in the model, as well as their interactions within specific temporal windows were independently and rigorously explored. I opted for this approach to avoid the theoretical demands of the proposed model, as well as experimental demands from previous studies. In addition to this, the principal parameter — *α* — was systemically explored, as well as the extent foveated systems require the presence of elφs to implement R. Specifically, for the simulation results reported below, the underlying architecture consists of two main layers – a “retinotopic field” and a “force field”. The retinotopic field is composed of hexagonal overlapping *RF_i_s* that tile the foveal, parafoveal, and peripheral extent of visual space with more *RF_i_*s tiling the foveal extent. Each *RF_i_* possessed corresponding eccentricity dependent elφs. Force fields perturbs *RF_i_s* from their equilibrium positions. This field is activated by the availability of “attentional-oculomotor signals” which align with early visually transient signals, the selection of a peripheral site, and later retinotopically modulated transient signals [12, 19]. The perturbation of *RF_i_s* produces spatiotemporal retinotopic trajectories which manifest in time-varying modulation of density at a given retinotopic location of visual space. I modeled each *RFi* as a bivariate gaussian kernel function **G,** Eq (**20**). To then obtain a probability density estimate for a retinotopic location, Eq. (**21)** was implemented, where *l*_1*i*_, *l*_2*i*_, denotes retinotopic locations from the bivariate distribution. *l* denoted the sampled retinotopic location whose density 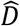 (or what I refer to as neural population density readouts) is being estimated over time. B denotes the bandwidth used, with G_B_ as a non-negative and symmetric function (∫ *G_B_*(*u*)*du* = 1) defined in bivariate terms as 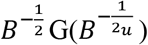.

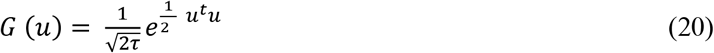

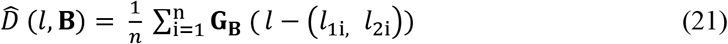

Population density readouts at the following retinotopic locations along the path of the impending saccadic eye movement (i.e., the radial axis) were sampled: outer foveal (*rad_fov–out_*), inner foveal/inner parafoveal (*rad_fov+para−in_*), outer parafoveal/inner peripheral (*rad_para+peri−in_*), and outer peripheral (*rad_peri–out_*) (Fig) **4**. It is important to highlight two very important points as it pertains to measurements at these retinotopic locations. One, results reported below are from when the centripetal force is fully turned on (*const*, *t*_*x*2_ < *t* < *t*_*x*3_) to when all forces including the centripetal force are turned off. The selection of this temporal window is consistent with previous remapping studies in foveated systems, where the organism is instructed to first acquire fixation and only after maintaining fixation allowed to preselect the peripheral target and then later execute a saccadic eye movement in the corresponding direction. Finally, because, visual sensitivity is lower at the more eccentric locations in visual space when compared to the more central locations during fixation (at t_0_), I controlled for known eccentricity effect [74] by dividing the estimated raw population density readouts at each sampled location with the mean of the first 110ms (i.e., the temporal window 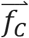 is in a steady state).

**Figure 4.**
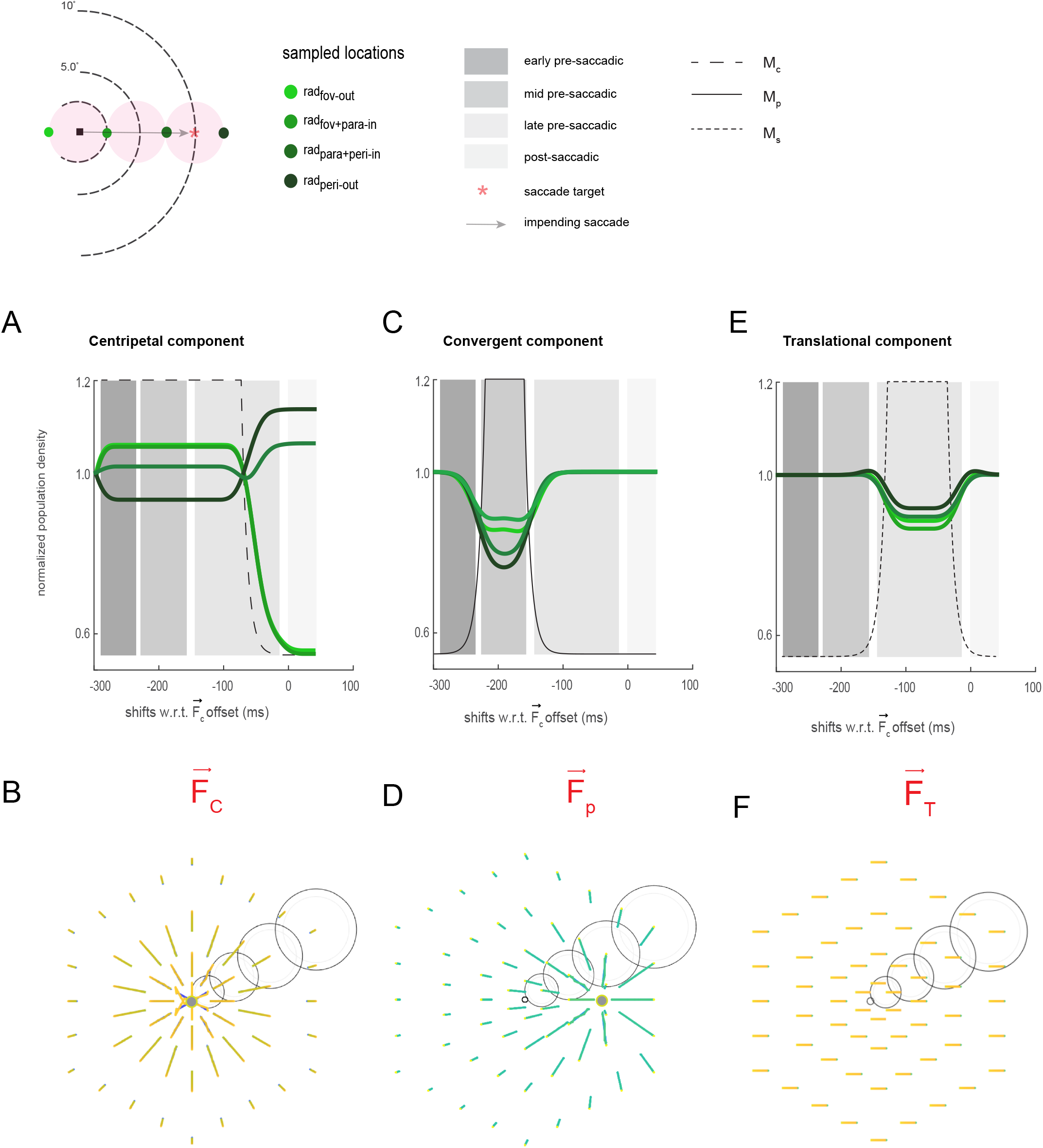
Single force simulations under an inverse force field. Simulation results (A) (C) and (E) at α = −1.6 in cases when RF_i_s are perturbed by a single external force constrained by elφs. Along x-axis represents the simulation time sampled at discrete increments of 1ms realigned with respect to the offset of the central force (i.e., modelled as the saccade onset). (B), (D) and (F) are corresponding spatiotemporal trajectories used to compute population density readouts.

#### Centripetal component

When modeled *RF_i_s* constrained by their elφs were perturbed by only a centripetal force Fig (**4A-B)**, I found distinguishable differences in population density levels between more foveal (rad_fov-out_ and rad_fov-in_) and more eccentric (rad_para+peri-in_ and rad_peri-out_) locations during the sustained period of this force. The decline of the centripetal force was accompanied by a clear hand-off in population density levels between the foveal and peripheral locations. I found an earlier and stronger increase in population density at the outer peripheral location (rad_peri–out_), followed by an increase in density levels at the inner peripheral location (rad_para+peri–in_).

#### Convergent component

Under the perturbation by a convergent force alone (Fig) **4C-D**, within a mid pre-saccadic window, I found a graded decline in population density levels across visual space. Under a strong inverse force field, the onset of a convergent force cause *RF_i_s* that support outer parafoveal and outer peripheral locations (rad_para+peri-in_ and rad_peri-out_) to allocate their resources towards the saccade target, while at the foveal and inner parafoveal locations (rad_fov-out_ and rad_fov-in_) neural resources remain largely stable. These results suggest that convergent signals are limited and do not provide neural advantages around a preselected peripheral site. On one level this result is deeply counterintuitive as one might expect convergent signals to produce effects that align with the convergent form of remapping. On another level, my finding is consistent with a more recent functional study [24] and a growing number of neural studies, which suggest that the functional and neural correlates of the convergent form of R fundamentally require an additional component [17, 18, 19].

#### Translational component

Similar to the simulations for the purely convergent case, I found a graded decline in population density levels across visual space due to a pure translational force applied during a late pre-saccadic window (Fig) **4E-F**. However, here, changes in population density levels favor the future center of gaze over the current center of gaze. These results align with the predictions proposed by early neurophysiological studies that the anticipatory transferring of resources from a cell’s current field to its future post-saccadic location is in part driven by oculomotor modulated translational signals [13–16, 66, 70]. On a functional level, this causes a relatively equal and transient decline in visual sensitivity across visual space [75].

#### Centripetal and convergent components

The combination of centripetal and convergent forces (Fig) **5A-B**. resulted in an amplification of the differences in population density levels along the radial axis during the application of the convergent force (compare with individual force simulations in (Fig) **5B**., while preserving the handoff noted earlier for the centripetal only simulation. Together with the centripetal simulation, these results predict the importance of the centripetal force in not only maintaining the appropriate level of resources at the current center of gaze during the sustained period of the force (i.e., at the time of fixation) but also ensuring the immediate availability of neural resources at the future center of gaze during the offset of the force (Fig) **4A**., (Fig) **5A**.

**Figure 5.**
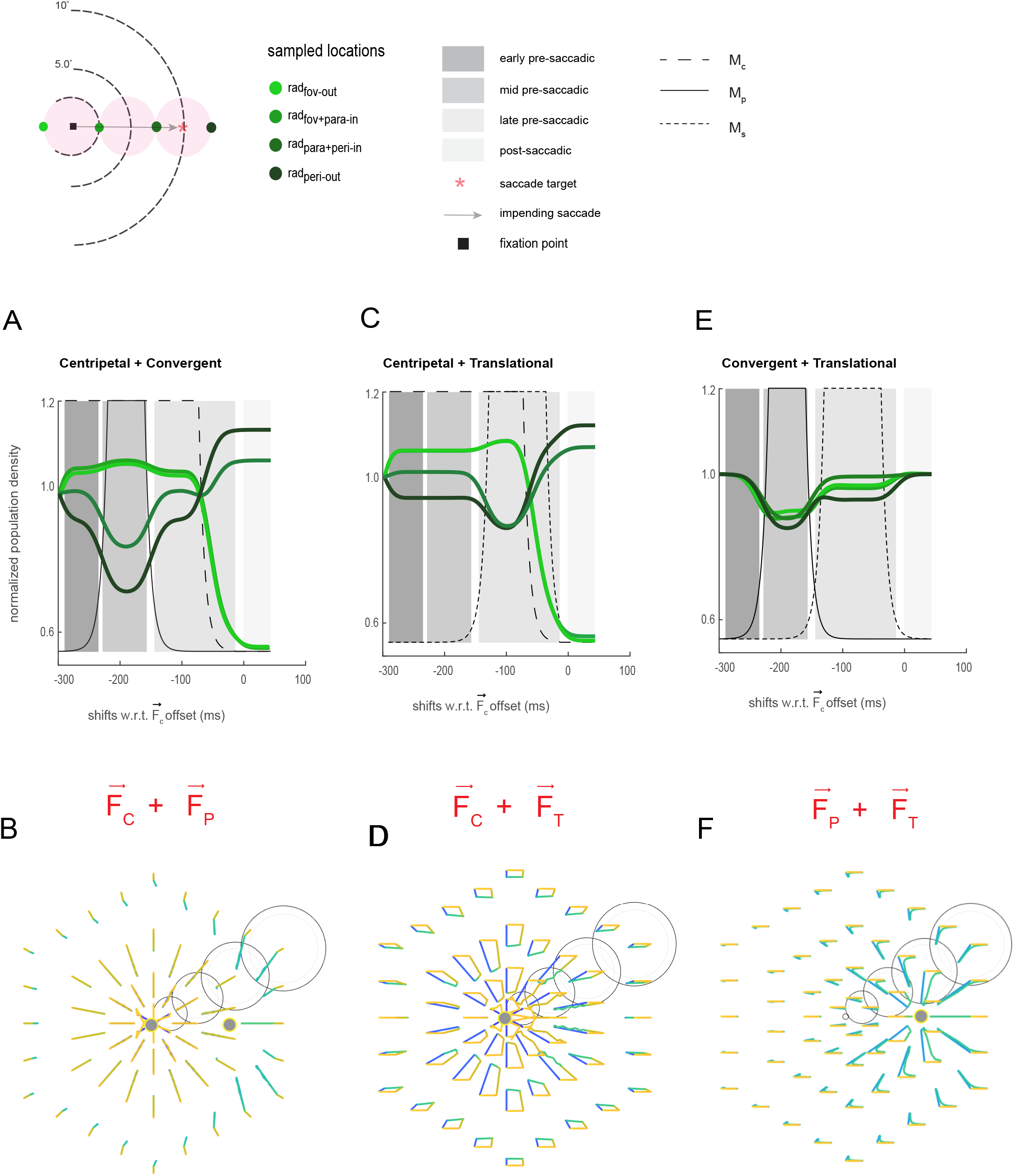
Two force simulations under an inverse force field. Simulation results (A) (C) and (E) at α = −1.6 in cases when when RF_i_s are perturbed by two external forces constrained by eccentricity dependent elφs. (B), (D) and (F) are corresponding spatiotemporal trajectories.

#### Centripetal and tranSlational components

The combination of the centripetal and translational forces resulted in an amplification of the divergence in density between the foveal and peripheral locations during the application of the translational force while retaining the foveal-peripheral handoff during the decline of the centripetal force (Fig) **5C-D**.

#### Convergent and translational components

With the combination of a convergent force and an increasing influence of a translational force, I found moderate asymmetric increases in population density levels at the rad_para+peri-in_ and rad_peri-out_ locations (Fig) **5E-F** when compared to the simulation by a convergent force alone (Fig) **4C-D**. No other simulation produced these changes in density levels in the periphery. On the neural domain, these transient increases in population density levels are consistent with a previous study that has shown transient increases in neural sensitivity in the frontal eye fields [18], which most likely corresponds to the preselection of the saccade target (i.e., a precursor of convergent remapping or the dynamics that initiates R in general) [12, 17, also see Fig 4B in 56]. On the functional domain, these results align with the prediction that increases in visual sensitivity just before saccade onset at a preselected peripheral site are not always symmetric [17, 57–59].

#### All components

The single and paired force simulations indeed capture specific components from the extant literature. These simulations however failed to conform to the idea that convergent signals alone or convergent along with translational signals provide significant neural advantages around the preselected peripheral site when compared to more distal locations (i.e., extents that are further away from the saccade target). This begs the question: does the combination of a centripetal, convergent, and translational force lead to such advantages? To directly investigate this question, I next conducted a simulation that included all three forces. To foreshadow these results, despite measurements from previous studies being tied to a specific retinotopic brain structure, the sampling of different retinotopic locations, as well as differences in quantification methods, results simulated by the proposed model revealed spatiotemporal dynamics that succinctly captured the essence of two very important empirical observations frequently cited in support of the translational [15] and convergent [19] forms of R.

Specifically, within an early pre-saccadic window I found a graded profile of neural density along the radial axis [50]: an immediate and sharp decline at the outer peripheral region (rad_peri–out_), a modest decline at the outer parafoveal region (rad_para+peri–in_) (Fig) **6A**, and sustained levels of neural density in the foveal (rad_fov–out_) and inner parafoveal (rad_fov+para–in_) regions (Fig) **7A**. This was followed by a modest rebound in density readout within a mid pre-saccadic window at the more eccentric locations (rad_peri–out_ and rad_para+peri–in_) from 240ms to 130ms before saccade onset, consistent with the model’s prediction related to the role of pure convergent signals. Within a late pre-saccadic window just before saccade onset, I found continued declines in density levels at locations near the current center of gaze (rad_fov–out_, rad_fov+para–in_). Within this same temporal window, I found a clear jump in neural density levels at a future post-saccadic location (rad_peri–out_) when compared to the more parafoveal extent (rad_para+peri–in_). To my surprise, these results qualitatively match the signatures reported by Sommer and Wurtz [15]. As Sommer and Wurtz’s results also predict, this jump in density followed a growing difference in density levels between the rad_peri–out_ location when compared to the more parafoveal location as the simulation approached saccade onset (Fig) **6A-B**.

**Figure 6.**
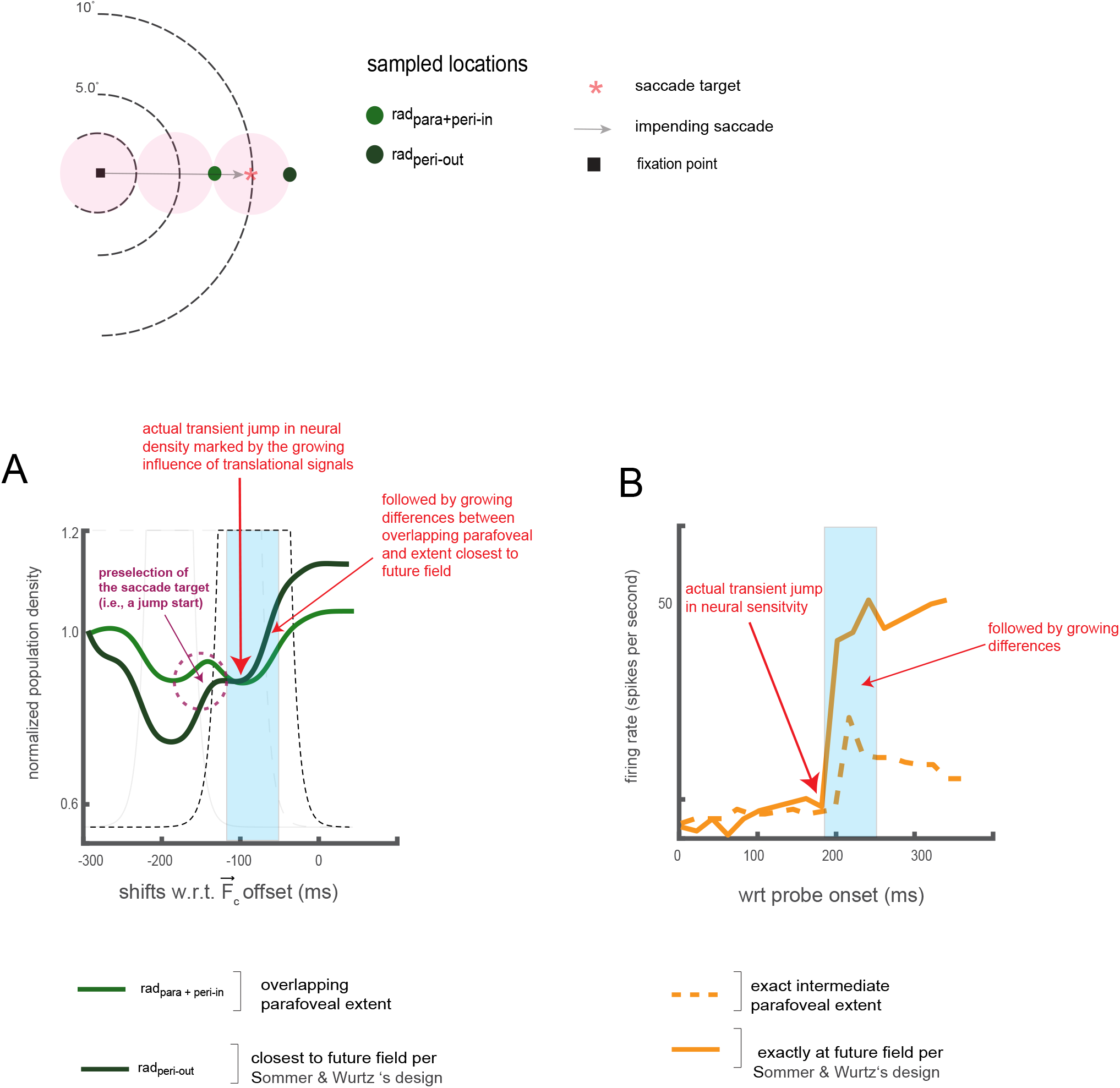
The combination of a centripetal, convergent and translational signals under an inverse force field qualitatively recapitulates translational effects reported by Sommer and Wurtz, 2006. Simulation results (A) Simulated population signatures constrained by eccentricity dependent elφs., under distinct inverse force fields of varying strengths. Along x-axis represents the simulation time sampled at discrete increments of 1ms (B) cartoon of results reported by Sommer and Wurtz, 2006, see figure 2c, right panel for exact plots.

**Figure 7.**
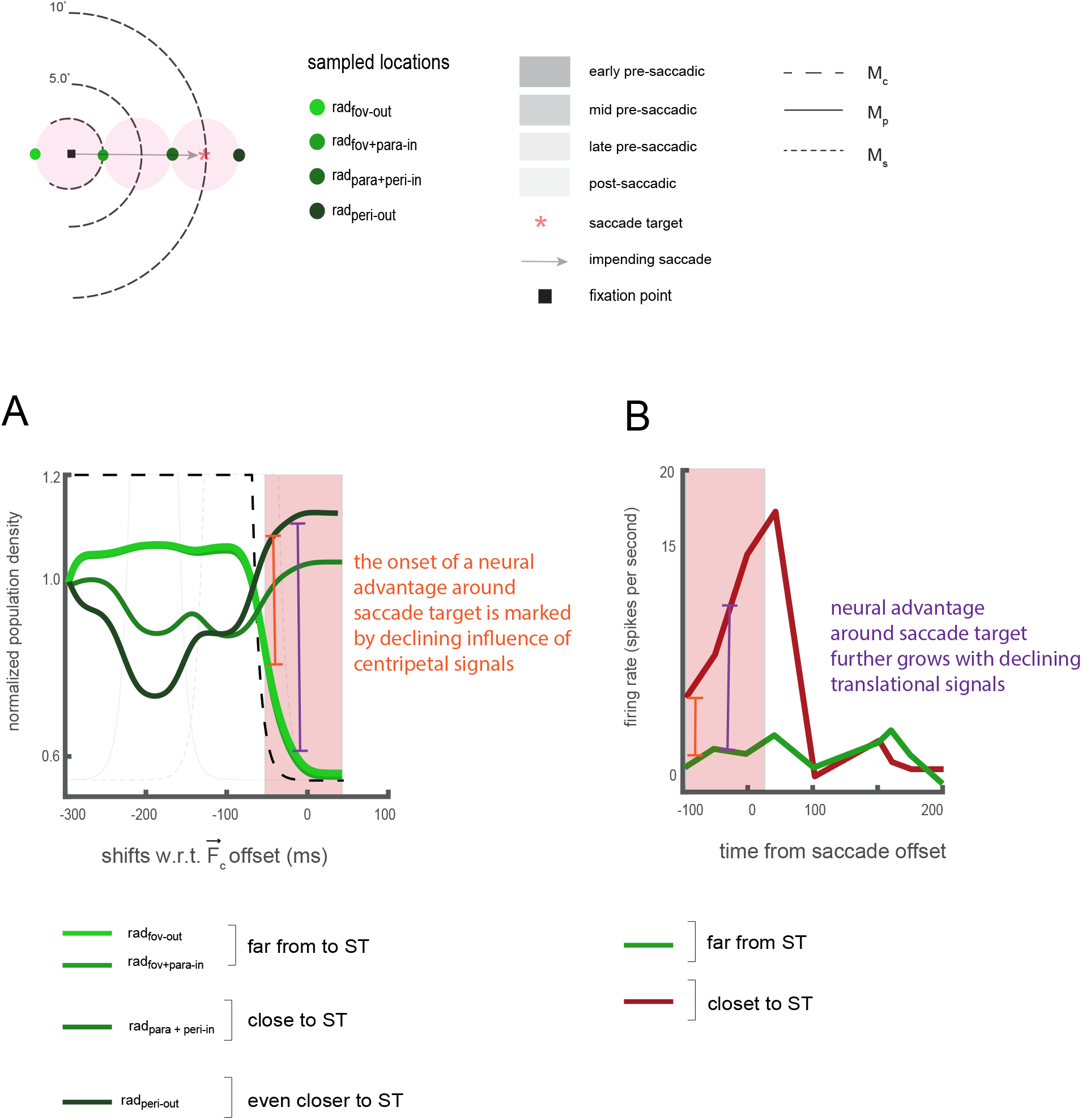
The combination of a centripetal, convergent and translational signals under an inverse force field recapitulates convergent effects reported by Hartmann et al, 2017. Simulation results (A) Simulated population signatures constrained by eccentricity dependent elφs under an inverse force field. Along x-axis represents the simulation time sampled at discrete increments of 1ms (B) cartoon version of results reported by Hartmann et al, 2017, see figure1B for the exact plot.

Finally, within a later pre-saccadic window along with an overlapping post-saccadic window, I found declines in post-saccadic density levels at locations furthest away from the saccade target (rad_fov–out_, rad_fov+para–in_), while the location close to the saccade target (rad_para+peri–in_) and an even closer extent (rad_peri–out_) (i.e., spatial extents around the simulated saccade target) demonstrated clear neural density advantages. Remarkably, these dynamics, which temporally align with the canonical order of pre-saccadic events [12], qualitatively match results reported by Hartmann and colleagues [19] (Fig) **7B**. These results for the first time reveal the importance of a declining centripetal force in implementing the convergent form of R, while the declining translational signals can further amplify neural advantages around a pre-selected peripheral site. Taken together, the recapitulation of the results by Sommer and Wurtz and Hartmann, and colleagues provides the long sought-after computational evidence that R is fundamentally a dynamic phenomenon [18, 21, 23, 24, 28, 29].

Additionally, the three-force simulation can also explain conflicting neural findings related to whether R is limited to the current and future post-saccadic locations. Specifically, classical neurophysiological studies have shown that *RF_i_s* respond to a salient cue placed within their future RF before the onset of an eye movement [13–16]. However, more recent results by Wang and colleagues [50] suggest that cells in LIP also become responsive to intermediate locations along the saccade trajectory. This is said to drive activity in favor of the current center of gaze that is eventually redirected (or remapped) towards the cell’s future post-saccadic location. The three-force simulation strongly suggests that graded changes in sensitivity towards the fovea occur uniquely within an early to mid pre-saccadic window (e.g., during early trainset signals), as predicted by Wang and colleagues. However, these graded changes in sensitivity cease within a late mid-saccadic and late pre-saccadic window just before saccade onset.

#### Exploring different a values

To assess the impact of the spatial profile of the force fields, I repeated the three-force simulations with different values of α, Fig (**8A-B**). With α = −1 (i.e., a moderate inverse force field) population density signatures at the more foveal locations are largely unchanged when compared to simulated results under a strong inverse force field where α is set to −1.6. More eccentric *RFis* are sensitized towards the current center of gaze but less so when compared to a strong inverse force field. To this end, within an early-mid pre-saccadic window I found declines in population density levels at the rad_para+peri-in_ and rad_peri-out_ locations. However, declines at the rad_peri-out_ location were not as large when compared to what I observed under a stronger inverse force field. Under a weak inverse force, where α is set to −0.6, intermediate parafoveal *RFis* are more sensitized towards the current of gaze. To this end, I observed larger declines in population density levels at the rad_para+peri-in_ location when compared to the simulations under a strong inverse force field. Biological systems are stochastic and sloppy, physical systems less so [76–79]. It is thus a certainty that between and within different biological systems, within different visual and movement contexts, α will vary. However, considering that stimulations under a moderate force field, as opposed to a weaker one in part, resembles signatures under a strong inverse force field, these results suggest, at least in foveated systems, that R likely obeys an inverse distance law (i.e., 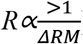).

**Figure 8.**
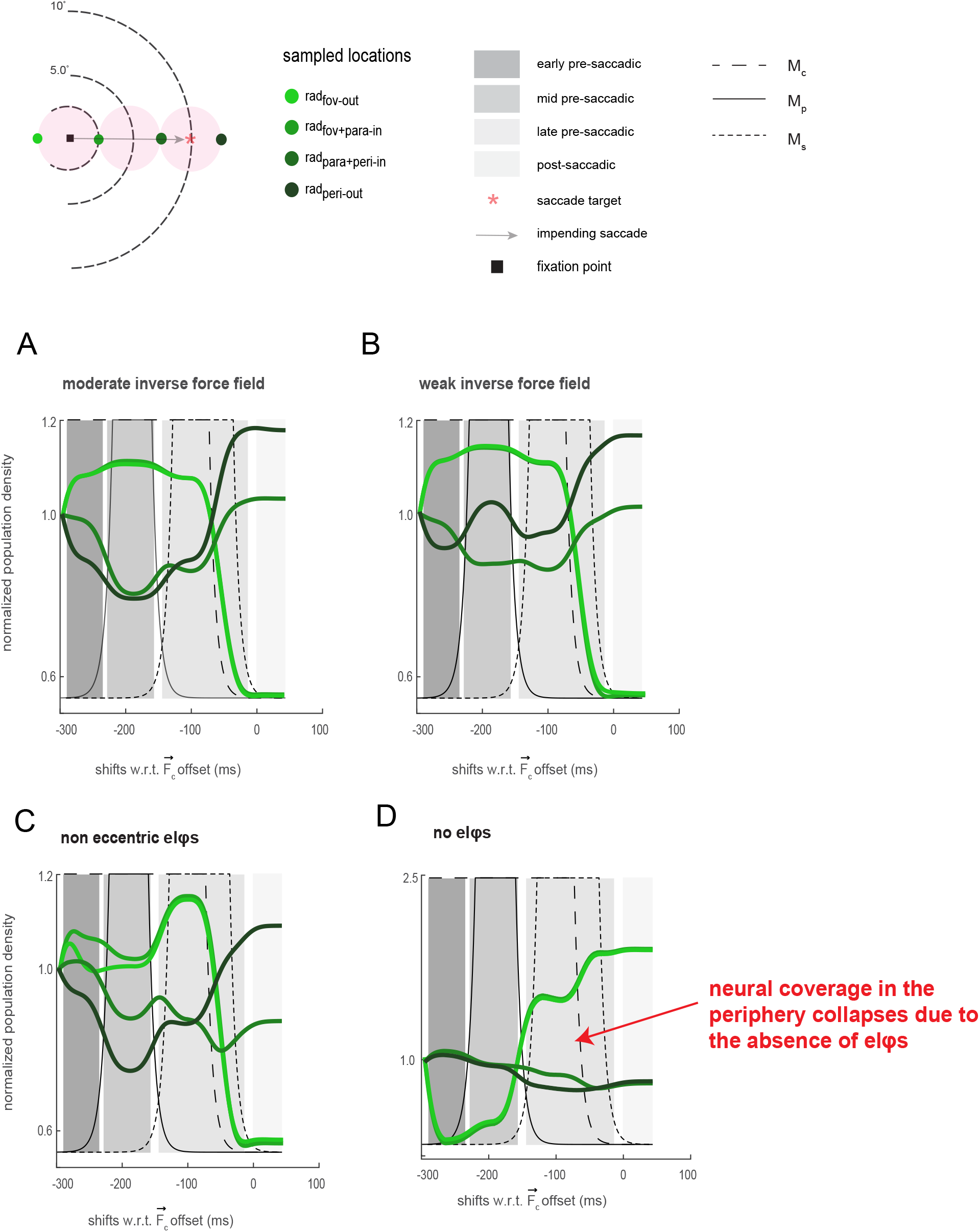
Additional three force simulations under an inverse force field. Simulated population density constrained by eccentricity dependent elφs, under distinct inverse force fields of varying strengths, (A) α= −1.0 and (B) α= −0.6. Simulated population density for the combination of three independent forces that are constrained by non-eccentricity dependent elφs (C) or unconstrained (D).

#### Exploring the architectural space

All of the results I have presented thus far were simulated with modeled *RF_i_s* constrained by eccentricity dependent elφs. To directly assess the role of elφs on pre and post-saccadic density readouts, I ran two additional simulations (both under a strong inverse force field, α = −1.6). For the first stimulation, ρβ was assumed with c set to a scalar greater than 1, while in the second stimulation pα was assumed with c set to 0. In the case of an eccentricity independent tilling of elφs, I found no appreciable differences in density levels when compared to the signatures I observed under eccentricity-dependent constraints during the early-mid pre-saccadic periods (Fig) **8C**; compare with (Fig) **6**–**7**. However, a different signature emerges with the introduction of a translational force. Specifically, I found that sustained density levels at the foveal regions increase in magnitude, while there is a marked decrease in density level at the more intermediate eccentric location. Eccentricity dependent elφs allow more eccentric *RF_i_s* a greater movement extent (i.e., more elasticity). Under a non-eccentricity-dependent configuration, however, eccentric *RF_i_s* are much more restricted in their ability to shift within the visual field. Consequently, once eccentric *RF_i_s* become sensitized towards the current center of gaze, they become restricted from reallocating these resources back towards the periphery. This results in an inappropriate level of sensitivity at the current center of gaze within a late pre-saccadic window as opposed to locations around the future center of gaze. Finally, for simulations where *RF_i_s* were unconstrained by their elφs, retinotopic organization simply collapses (Fig) **8D.** The centripetal force results in an over-compression towards the current center of gaze, with the peripheral region experiencing radical declines in sensitivity. Further, despite the onset of the convergent force (e.g., the looming presence of a predator, nearby car, etc.) *RF_i_s* are largely desensitized to the presence of these forces [80–82]. This result points to the importance of elφs in preventing any radical and unsustainable forms of remapping while ensuring that retinotopic organization is always maintained.

### B. PSYCHOPHYSICS

To now assess the biological plausibility of the model and its predictive power directly, I conducted three psychophysics experiments in foveated systems (human subjects). Specifically, I assessed changes in visual sensitivity across visual space using a cued saccade task (Fig) **9**. To do this, I carefully mapped the geometry of visual space with the intent to examine transient consequences of predictive remapping on visual sensitivity changes at precise points along the saccade trajectory (‘radial axis’) matching the locations I simulated using the proposed model. In addition, symmetric points orthogonal to the saccade trajectory (‘tangential axis’) were also measured. Visual sensitivity measurements were obtained while human subjects fixated, selected a target on the peripheral retina, and planned, and executed a saccadic eye movement toward the peripheral target. Specifically, radial points at foveal (rad_fov–out_, rad_fov–in_), parafoveal (rad_para–in_, rad_para–out_) and peripheral (rad_peri–in_, rad_peri–out_) locations, and tangential points symmetric to one another with respect to the radial axis at foveal (tan_fov–ccw_, tan_fov–cw_), parafoveal (tanp_ara–ccw_, tanp_ara–cw_) and peripheral (tan_peri–ccw_, tan_peri–cw_) locations were probed, before and after the central movement cue (Fig) **9A**. In each trial, a low contrast probe was flashed at one of these locations (chosen at random) for 20ms with 75% probability; in 25% of trials (control trials), no probe was flashed. The contrast of the probe stimuli was chosen (independently for each subject and for each spatial location cluster; see Methods) such that the detection probability was at 50% in the absence of eye movements. The probes were presented at a random time from 600ms before to 360ms after the central movement cue. This experimental paradigm was attractable for two reasons. One, most remapping studies typically sample locations in space favorable to one form of remapping over another. However with this dense spatial sampling of visual space, this allowed me to assess the possible primary role the centripetal, convergent, and translational components play in modulating visual sensitivity within distinct regions of visual space. Finally, measuring visual sensitivity along and around the entire saccade trajectory, within early pre-saccadic (−300ms to - 250ms with respect to saccade onset), mid-pre-saccadic (−250ms to −150ms), late pre-saccadic (−150ms to - 5ms) and post-saccadic (−5ms to +150ms) windows was critical in assessing the consequences of R on visual sensitivity within a larger pre and post saccadic window (Fig) **9B**.

**Figure 9.**
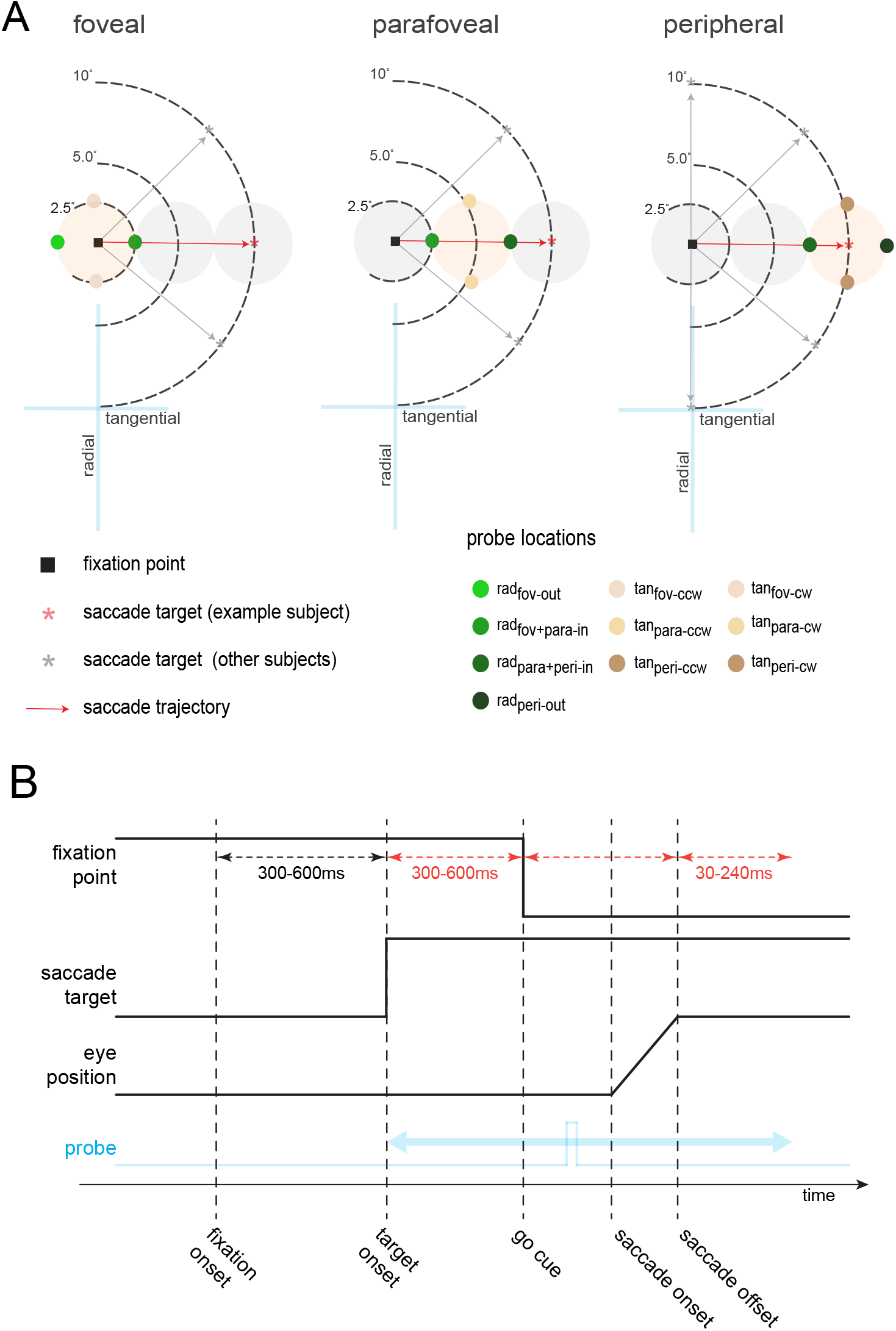
Cued probe-detection task. (A) Spatial locations tested across experiments. Colored circles show the locations at which a low-contrast probe was flashed as subjects planned and prepared a saccade (black arrow) towards the periphery. Probe locations are shown only for horizontal saccades to the right. For other saccade target locations, the probe locations were appropriately positioned along and around the path of a saccade. In the foveal and parafoveal experiments, a saccade target (gray or red asterisk) was presented at azimuth angles of 0°, 45°, 315° (each subject had a different angle). In the peripheral experiment the saccade target was presented either at azimuth angles of 0°, 45°, 90°, 270°, 315° (each subject had a different angle). (B) Temporal sequence of an example trial. The start of a trial began with the appearance of a fixation point, followed by a time-varying presentation of the saccade target. A low-contrast probe was flashed either before the onset of the central movement cue (the pre-saccadic condition), or after (the post-saccadic condition).

Subjects reported whether they were able to detect the flashed probe using a push button. The control (no probe) trials allowed me to assess the incidences of false alarms. Low false alarm rates of 1.2%, 1.4%, and 1% (along the radial, tangential counterclockwise, and clockwise axes respectively) gave me high confidence about the subjects’ visual sensitivity reports. I collected ~ 21,000 trials, with a minimum of 1,500 trials per subject. Control trials were excluded from the main analyses. I calculated the normalized average visual sensitivity of eleven subjects as a function of flashed probe times relative to saccade onset (See Supplementary (Fig) **2**, for control readouts between groups for the parafoveal and peripheral experiments). Corresponding error estimates were obtained using a 20-fold jackknife procedure in which the sensitivity was estimated from 95% of the data (See Supplementary **Methods** for experimental protocol and details regarding behavioral analyses).

Within an early pre-saccadic window (−300ms to −250ms), a graded visual sensitivity profile along the radial axis was observed, consistent the results reported by Wang and colleagues [50], and simulated results (Fig) **6**–**7**, **10A**. Within a mid pre-saccadic window (−250ms to −150ms) before saccade onset a modest rebound (or a jump start) in visual sensitivity at rad_peri–out_ and rad_para+peri–in_ locations was observed, consistent with the predicted role of convergent signals and simulated results. Continued declines in visual sensitivity at locations near the *rad_fov–out_*, *rad_fov+para–in_* were observed and consistent with classical neurophysiological findings [13–16], as well as simulation results. Declines in sensitivity at the rad_fov–out_, rad_fov+para–in_ location were observed along with a jump in late pre—saccadic (−150ms to −5ms) sensitivity at the radperi-out location. Finally, within a post-saccadic window continual declines in post-saccadic sensitivity at the rad_fov–out_, radfov–in location were observed along with a sharp and immediate increase in post-saccadic (−5ms to +150ms) sensitivity at the rad_peri–out_, rad_para+peri–in_ locations. These sensitivity signatures not only align with simulated results but are consistent with previous functional and neurophysiological findings [32, 24, 17, 18, 19].

**Figure 10.**
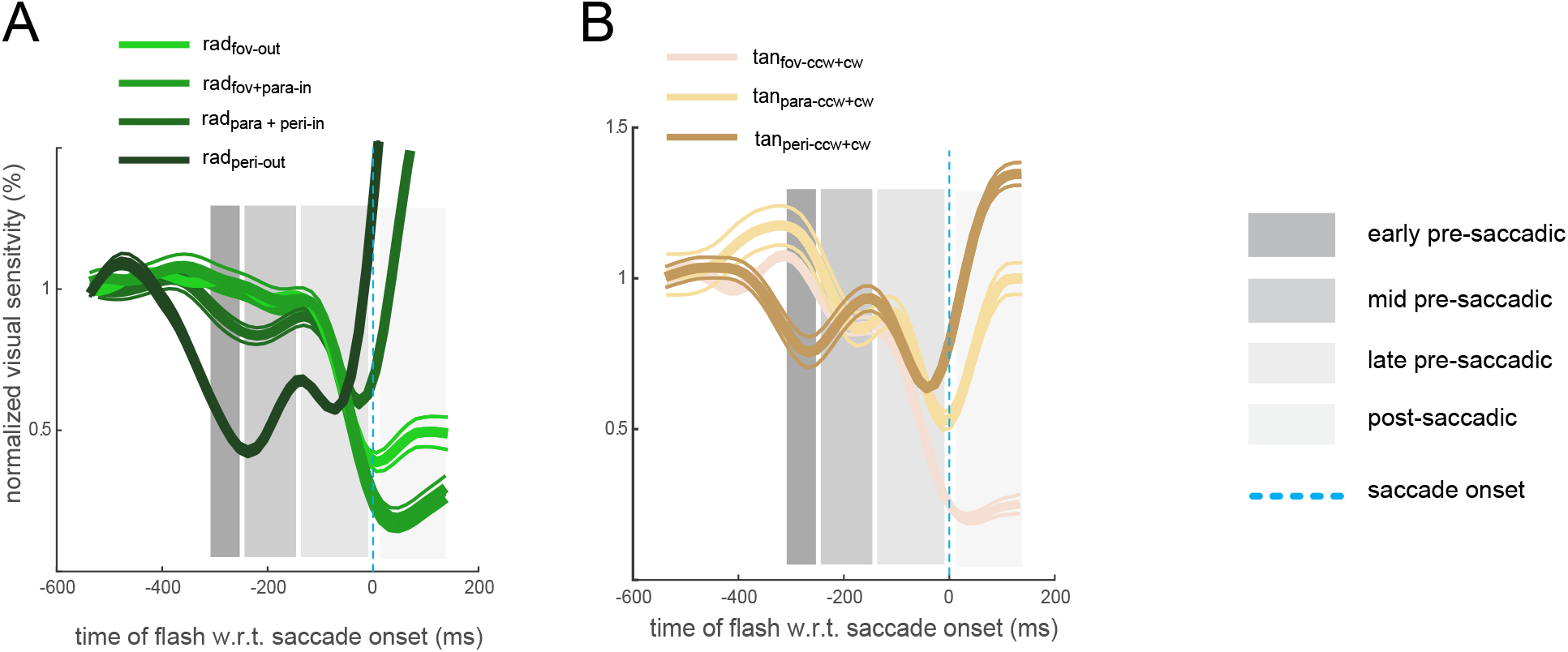
Sensitivity functions across visual space within distinct pre and post saccadic windows.Thick solid lines represent normalized changes in pre and post-saccadic sensitivity as a function of flashed probe times relative to saccade onset along (A) and around (B) the saccade trajectory (n=11). The thin lines represent corresponding error estimates obtained using a 20-fold jackknife procedure in which the sensitivity was estimated from 95% of the data.

Observations of visual sensitivity changes along the tangential axes largely mirrored those along the radial axis in an eccentricity dependent fashion (Fig) **10B**. I found a decrease in sensitivity at the tan_peri–ccw+cw_ location within early pre-saccadic window. This was accompanied by a modest increase in mid pre-saccadic sensitivity in the parafoveal region (tan_para–ccw+cw_) or sustained levels of sensitivity in the foveal (tan_fov–ccw+cw_) region. Similar to the radial axis, I observed a rebound in late pre-saccadic sensitivity (−50ms from saccade onset) at the most eccentric location (tan_peri–ccw+cw_) that was followed later and less marked rebound at the parafoveal location (tan_para–ccw+cw_). A continued decline in post-saccadic sensitivity at the foveal location was accompanied by rapid increases first at the peripheral locations followed by the parafoveal location.

Quite convincingly, the combination of three independent forces largely predicts sensitivity readouts along the radial and tangential axes, except for the tan_peri–ccw_ location during an early and mid-pre-saccadic window. These results point to the biological plausibility and generalizability of the proposed model, Supplementary Fig (**3**). On one level, these results contradict a set of functional studies that support the translational form of R [e.g., 26 27], all of which suggest that there is no spreading of neural resources around the pre-selected saccade target. On another level, the principal suggestion from these functional studies that pure convergent signals cannot provide the type of neural advantages around the saccade target that previous neural studies have reported holds (Fig) **4C-D**. To conclude, increases in post-saccadic sensitivity at the locations around the saccade target (tan_peri–ccw+cw_, *rad_peri–out_*, *rad_para+peri–in_*) compared to the more distal locations (tan_fov–ccw+cw_ rad_fov–out_, rad_fov+para–in_) provide the long-sought after functional evidence in support of the convergent form of R. Mechanistically speaking, the onset of this neural advantage is primarily due to the presence of declining centripetal signals with translational signals playing more of a secondary role.

### C. RHYTHMICITY

Having demonstrated that R is a dynamics phenomenon that includes the newly discovered centripetal component along with convergent and translational components, I next wanted to assess a core assumption of the model, that is, early visually transient and feedback transient signals are uniquely supported by delta rhythmicity (2 to 4Hz). Indeed, this rhythmic band is known to predict attentional shifts before the onset of a saccadic eye movement [65].

I hypothesized that attentional signals are highly selective towards the current and future center of gaze, while at intermediate parafoveal locations retinal image displacement signals blunt attentional resources that would otherwise persist at these locations. To test this, I assessed the spectral signature underlying the observed changes in sensitivity [83]. The logic behind this assessment is as follows: if these changes in the center of gaze and the peripheral region where the saccade target resides are a consequence of attentional shifts, then the spectral signature associated with changes in sensitivity at these locations should be statistically significant and resemble, if not match, known rhythmic patterns of attentional shifts during fixation and the selection of the peripheral target. Conversely, in the parafoveal region observed rhythmic patterns should not be associated with attentional shifts. To this end, if the spectral architecture I find fails to recapitulate these known rhythmic patterns in the foveal and peripheral regions or I statistically significant in the parafoveal regions, then sensitivity changes at the center of gaze and peripheral region cannot be directly attributed to attentional shifts.

Transforming detrended times series sensitivity data (Supplementary Fig (4) into the frequency domain, I found that visual sensitivity around the current center of gaze (“foveal”) locations (rad_fov–out_, rad_fov+para–in_) exhibited rhythmicity within a delta spectral window at ~2 Hz, except for the tan_fov–ccw+cw_ location which exhibited rhythmicity that bordered the delta and theta windows at ~3 Hz (Fig) 11A. Spectral differences calculated using location shuffled data (n=1500) versus the non-shuffled data at these locations were statistically significant (two-tailed paired-sample t-test, p=0.0018, p=0.0236, p=0.0018). These results not only align with my but are consistent with the idea that delta rhythms at ~3Hz actively mediate pre-saccadic attentional shifts at suprathreshold levels [65, 84–86]. At the intermediate (“parafoveal”) locations (tan_para–ccw+cw_), I found that sensitivity exhibited delta-band rhythmicity at ~2Hz (Fig) **11B**. However, the spectral differences between the shuffled and non-shuffled data sets were not significant (p=0.1738). Finally, peripheral locations (Fig) 1**1C**. around the future center of gaze (tan_peri–ccw+cw_, rad_para+peri–in_, rad_peri–out_) all exhibited rhythmicity at 2-3Hz which like in the foveal region, were statistically significantly different from location-shuffled data (p= 0.0062, p=0.0071, p= 0.0207 respectively).

**Figure 11.**
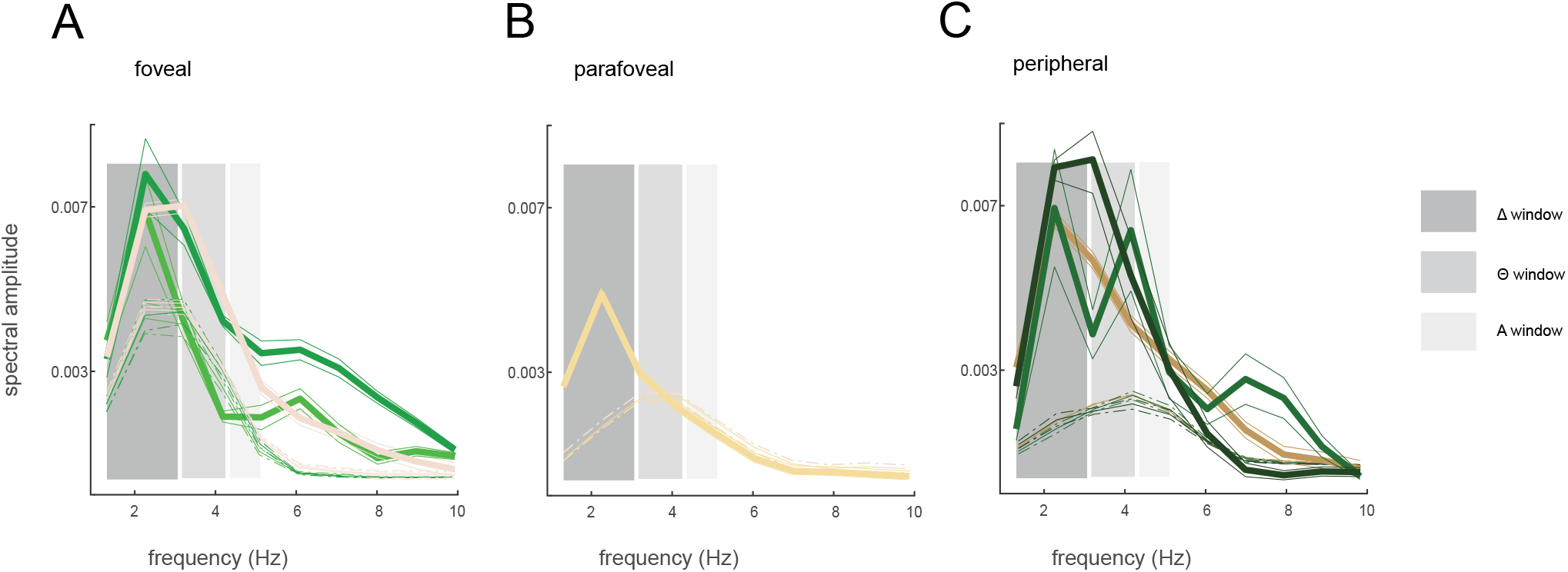
Frequency domain representation of detrended sensitivity functions within distinct spectral windows. Frequency domain representation of detrended visual sensitivity, unshuffled (solid lines) and shuffled (dotted lines, n=1500) in the fovea (A), parafovea (B) and periphery (C). The solid and dotted thinner solid lines represent the error estimates calculated across subjects.

Taken together, these results suggest that attentional signals support changes in visual sensitivity in task-relevant regions of visual space around the current [55, 61]and future centers of gaze [24, 32]. However, at the intermediate parafoveal locations, the region of visual space susceptible to the incessant retinal disruptions saccades necessarily accompany, these changes in visual sensitivity are not strongly mediated by attentional signals. Concerning the proposed model, these results strongly suggest R may be driven by “attentional oculomotor signals” that are actively being mediated by delta rhythmicity.

## IV. DISCUSSION

Predictive remapping is a phenomenon actuated by cells across different retinotopic brain structures. Indeed, this phenomenon has been observed across different sensory modalities [13–19, 87–88] and across different foveated biological systems which include humans [89] and rhesus macaques. Here I was motivated by what is arguably the most puzzling question in the field of predictive remapping research. In 1990 Goldberg and Bruce recapitulated translational effects observed in the superior colliculus by Mays and Sparks ten years earlier [90, 33]. They found that before the execution of a saccade, the receptive fields of cells in the frontal eye fields predictively shift their spatial extent beyond their classical extent towards their future fields (translational remapping). These neural effects were reproduced across several retinotopic brain structures including the lateral intraparietal area and extrastriate visual areas. About two decades later, recapitulating in part the results of Tolias and colleagues [17], Zirnsak et al. demonstrated that the receptive fields of cells in the frontal eye fields primarily converge around the attention-selected peripheral region of interest which included the spatial extent of the saccade target (convergent remapping) [18].

I revisited these contradictory findings by introducing the Newtonian model of predictive remapping in which I proposed that the underlying pre-saccadic attentional and oculomotor signals manifest as temporally overlapping forces that act on retinotopic brain structures. Next, I systematically assessed the transient consequences of predictive remapping on visual sensitivity along and around the path of a saccade. I show that, contrary to the dominant spatial account, predictive remapping is neither a purely convergent nor a translational phenomenon but rather one which must include a translational, convergent, and centripetal component. I demonstrate, contrary to the dominant restrictive account, predictive remapping is not limited to the current and future post-saccadic locations but includes intermediate parafoveal locations along the saccade trajectory within the early to mid-pre-saccadic window [50]. This transiently shifts neural resources towards the current center of gaze that is eventually remapped towards the future center of gaze. Finally, I show that, contrary to the dominant temporal account of predictive remapping, centripetal shifts towards the fovea precede and overlap with convergent shifts towards the peripheral region of interest, while translational shifts parallel to the saccade trajectory occur later in time and overlap with convergent shifts.

This study makes four principal contributions to a deeper understanding of predictive remapping. First, recipient retinotopic brain structures receive temporally overlapping inputs that align with a canonical order of pre-saccadic events. Second, the neural computations that underlie predictive remapping obey an inverse distance law. Third, during active sensing (when not in a state of equilibrium) *RFi* very likely possesses putative transient eccentricity dependent elastic fields (elφs). It is likely that elφs are self-generated by the visual system and allow these receptive fields to undergo a remarkable degree of elasticity beyond their classical spatial extent. Simulation results show, in the absence of these elφs, orderly perception of the visual world would be dramatically distorted, as retinotopic cells would over-commit to certain loci in visual space at the expense of others which would compromise the ability to detect changes that occur on the peripheral retina [80–82]. Finally, this study strongly suggests that the immediate availability of neural resources after the execution of a saccadic eye movement is uniquely mediated by centripetal signals directed toward the current center of gaze. Indeed, without centripetal signals or elφs, attentional resources that are needed at the approximated location of the target would be significantly delayed and in the extreme case non-existent (Fig) **8D.**

In conclusion, I propose a simple but powerful mathematical abstraction that was implemented by the proposed model [91]. This novel framework uncovers a fundamental law and the functional architecture that mediates predictive remapping, which had eluded the field for decades. It not only offers critical insights that will inform future investigations of this phenomenon, specifically those concerned with uncovering the functional and neurobiological basis of elφs, but a powerful tool that experimentalists can now use to generate novel predictions across different types of biological systems.

## V. SUPPLEMENTARY MATERIAL

### A. REMAPPING EXPERIMENTS

#### Consenting human subjects

Eight subjects, all with corrected-to-normal visual acuity, provided written and informed consent before the study. Each subject, with the exception of two individuals (both authors), was compensated 20 USD per hour for their participation. The protocol for this study, the collection, and storage of the data was approved by the Yale Ethic Review board and was in accordance with the Declaration of Helsinki.

#### Viewing distance

Subjects sat comfortably with their chin and forehead placed against a custom-built head support apparatus consisting of an adjustable chin rest, forehead support and a bite bar holder. The center of the experimental display was placed at a distance of 57cm from the eye. In order to ensure tight control over viewing distance and eye-position, subjects placed a bite bar (personalized deep-impression dental bite-bar) inside their mouth. The bite bar was then affixed to the head support apparatus for the duration of the experimental session.

#### Stimulus and presentation

Visual stimuli were presented on a gamma corrected LCD monitor using custom-designed Window software (Picto). All stimuli presented in the control and main experimental sessions, with the exception of the achromatic Gabor probes (0.5 degrees of visual angle [dva], 0° orientation, π/2 phase), were constructed in Picto. The display was driven by a NVIDIA graphics card with a color resolution of 8 bits per channel. Its spatial resolution was 1400 × 1050 pixels with a refresh rate of 85 Hz (11.76 ms per frame), and an average mean luminance of 38 cd/m2.

#### Eye-movement Tracking

The right eye (the dominant eye for all subjects) was tracked using an infrared video-based eye-tracker sampled at 1kHz (I-Scan Inc., Woburn, MA). At the beginning of each experimental session, I performed a custom-developed 9-point eye-calibration procedure in Picto. This procedure allowed for the correction of any drift in gaze at the onset of a trial. Overall, the gaze error within subjects varied from 0.25° to 1.0° with an average gaze error of 0.45° across subjects.

#### Contrast Sensitivity Function measurement

Prior to the main task, I measured the Contrast Sensitivity Function (CSF) at an eccentricity corresponding to the mean location at which probes were presented for the main task. CSF was measured using a set of Gabor stimuli contrasted against a grey background with 21 contrast levels ranging from 1 to 55%. At the onset of each trial, subjects were instructed to acquire and maintain fixation for at least 1000ms. Trails were aborted if the eye-position deviated from fixation by more than 1 dva. A Gabor stimulus with a randomly selected contrast level was then flashed for 20ms. Subjects had to indicate with a button-press whether they were able to detect the stimulus. Each contrast level was presented 10 times, along with a non-probe condition, which controlled for any potential false alarms. After the conclusion of this session, a logistic function was used to estimate a psychometric curve fitted to the data to compute a CSF. The contrast at which the subject could detect the stimuli with 50% accuracy was chosen as the probe contrast level for the main task. The CSF measurement sessions lasted approximately 70 minutes.

#### Cued saccade task

In the main experiment, subjects performed a cued saccade task (Fig) **10**. Before the commencement of an experimental session, each subject took ~15 minutes to adapt to the scotopic condition in the experimental room. At this time, a 9-point eye-position calibration procedure was performed (see above). Subjects initiated a trial by maintain gaze on a central fixation dot (subtending 0.5 dva in diameter) for 300ms. If the eye deviated from this dot by more than 1 dva, the trial was aborted. After a variable delay period of 0-300ms a saccade target (white circle) appeared at a 10° eccentric location. Subjects were required to maintain fixation for another 300-600ms, at the end of which the fixation dot was extinguished, which served as the central movement cue to make a saccade towards the saccade target (Go_onset_). The saccade target remained on the display for another 500ms after the central movement cue. A 2-dva radius tolerance window around the saccade target was used to detect whether the saccade landed on the target within this 500ms window. Subjects were provided visual feedback about the outcome of the trial by changing the colour of the saccade target to green for a correct saccade landing within the tolerance window or red for saccades landing outside the tolerance window. On 25% of the trials a Gabor probe (contrast set to 50% detection rate from the CSF measurement), was flashed for 20ms at a random time during the 300-600ms window between the onset of the saccade target and the central movement cue. On 50% of the trials, the probe was presented at a random time between 0-340ms after the central movement cue. To control for any false alarms no probe was flashed in the remaining 25% of trials. These three conditions were randomly interleaved across trials. Subjects indicated whether they detected the probe by a button press. Consecutive trials were separated by a 1000ms inter-trial interval. The spatial locations of the flashed probes depended on the experimental conditions outlined below. Each experimental session lasted about 70 minutes with a 10-minute break in the middle. Subjects took about 3 weeks to complete the main experiment.

#### Fovealprobes condition

3 subjects (all female, Mage= 22, SD =1.7) were recruited for the foveal condition. Each subject was completely naïve to the aim of the experiment. Saccade targets appeared at an eccentricity of 10° and at azimuth angles of 0° (for s1), 45° (for s2) or 315° (for s3). In the event a flashed probe was displayed (on 75% of trials), this occurred with equal probability at one of 4 isotropic locations which were 2.5 dva from the fixation dot. Two of these locations were along the radial axis (the axis collinear with the fovea): farther away from fixation (rad_fov–out_), and between the fixation dot and the saccade target (rad_fov–in_). The other two locations were along the orthogonal (tangential) axis: counterclockwise (tan_fov–ccw_) or clockwise (tan_fov–cw_).

#### Parafoveal probes condition

The same group of subjects who participated in the foveal condition took part in the parafoveal condition. The location of the saccade targets was the same as in the foveal condition. Here, the probes were presented around the midpoint (at eccentricity of 5°) between the fixation dot and the location of the saccade target (i.e., at the parafoveal retinotopic location mid-way along the saccade trajectory). Each flashed probe appeared 2.5° dva from this parafoveal midpoint along the radial-tangential axes. Specifically, a probe could either be flashed at the most inner radial parafoveal location (rad_para–in_) (i.e., location closest to the fovea, which overlaps with the rad_fov–in_ probe location in the fovea experiment), the least inner radial parafoveal location (rad_para–out_) (i.e., location furthest away from the fovea) or along the tangential axis through the parafoveal midpoint either counterclockwise (tan_para–ccw_) or clockwise (tan_para–cw_).

#### Peripheral probes condition

5 subjects (2 Males, 3 Females, Mage= 25, SD =2.7) were recruited for this condition. With the exception of subject EL(s_EL_, the author), each subject was completely naïve to the aim of the experiment. Depending on the subject, the saccade target was presented at an eccentricity of 10° with azimuth angles of 270° (for s1), 315° (for s2), 0° (for s3), 45° (for s4), 90° (for s_EL_). In this condition a probe was flashed 2.5° dva from the saccade target along the radial-tangential axis. Specifically, a flashed probe appeared at either the inner radial location (rad_peri–in_, the same location the rad_para–out_ probe was flashed), the outer radial location (rad_peri–out_), or along the tangential axis either counterclockwise (tan_peri–ccw_) or clockwise (tan_peri–cw_) with respect to the saccade target.

### B. DATA ANALYSIS

#### Valid trials

Eye-position and push-button responses obtained from each subject were recorded at 1kHz and stored for further analyses. A trial in which the subject’s eye-position landed within the 2-dva tolerance window around the saccade target and within 500ms of the central movement cue was considered a valid trial. Across all subjects I collected a total of 5826 valid trials for the foveal condition, 5579 valid trials for the parafoveal condition, and 9473 valid trials for the peripheral condition.

#### Saccade onset and offset estimation

For each valid trial, I estimated the onset and offset of the saccade using a “displacement method”. Specifically, I calculated the variance in eye-position before the central movement cue and after the eye landed on the saccade target. I then estimated the lower and upper bounds of the 95% confidence intervals which accounted for any possible spurious movement which occurred before and after the saccade. The time points at which the eye-position deviated from these bounds were used to estimate the start (S_onset_) and end (S_offset_) of the saccade. S_onset_ and S_offset_ estimated as further verified using Cluster Fix, a software which performs k-means clustering on the following parameters: angular velocity, acceleration, distance and velocity in order to classify a rapid eye movement [92].

#### Excluding trial with corrective saccades

I eliminated trials with corrective saccades (i.e., secondary saccades which compensated for under- or overshoots in the primary saccade), since such trials could potentially be associated with non-canonical sensitivity dynamics. Analysis revealed that such trials always had saccade durations (including both the primary and secondary components) greater than 50ms. I therefore eliminated trials with total saccade duration greater than 50ms, and also visually verified that such trials contained a corrective component. In the foveal condition, of the 5826 valid trials, 17 trials were eliminated for subsequent analyses (no-probe =6; rad_out–fov_=3; rad_fov–in_=3; tan_fov–ccw_=2; tan_fov–ccw_ =3). Similarly, 6 out of 5579 valid trials were eliminated in the parafoveal condition (no-probe =1; rad_para–out_=0; rad_para–in_=3; tan_para–ccw_=1; tan_para–cw_ =1), and 12 out of 9473 valid trials were eliminated in the peripheral condition (no-probe =1; rad_peri–out_=3; rad_peri–in_=3; tan_peri–ccw_=1; tan_peri–cw_ =4).

#### False alarm rate

I calculated the false alarm as the probability of a button press in trials in which no probe was presented. Subjects performed the task with very low false alarm rates in all experimental conditions: 2%, 1% and 1% respectively in the foveal, parafoveal and peripheral conditions.

#### Raw fluctuations in sensitivity

To estimate the visual sensitivity at different retinotopic locations during a rapid eye moment, I isolated the valid trials which included a flashed probe at one of the four spatial locations in the foveal, parafoveal and peripheral conditions. I first realigned the trial data to the saccade onset time (time zero) and calculated the probe presentation time with respect to saccade onset. I then calculated sensitivity, separately for the four spatial locations, by using a 25ms sliding window that was moved in 15ms increments. For each time window I calculated the fraction of trials with button pushes over the number of trials in which a probe was presented. To obtain a robust estimate of each subject’s sensitivity, I employed a 20-fold jackknife procedure in which the sensitivity was estimated from 95% of the data. This was repeated 20 times, each time leaving out 5% of the data, giving us a mean and error estimate of the sensitivity for each spatial location in each experimental condition (rad_fov–out_, rad_fov–in_, rad_fov–tcc_, rad_fov–tcw_ probes in the foveal experiment; rad_para–out_, rad_para–in_, rad_para–tcc_, rad_para–tcw_ probes in the parafoveal experiment, and the rad_peri–out_, rad_peri–in_, rad_peri–tcc_, rad_peri–tcw_ probes in the peripheral experiment).

Keeping in mind that in the foveal and parafoveal experiment, flashed probes at the rad_fov–in_ and the rad_para-in_ locations, and in the parafoveal and peripheral experiment, flashed probes at the rad_para–out_ and rad_peri–in_ locations were subtended at the same retinotopic location, I further combined the jackknife sensitivity data at these locations. In the same vein, symmetric points along the tangential location in the foveal, parafoveal and peripheral regions were also combined. Consequently, I obtained a total of seven radial and tangential sensitivity functions: four along the radial axis (rad_fov–out_,rad_fov–in+radpara-in_, rad_para–out+radperi–in_, rad_peri–out_), three along the tangential axes (tan_fov–ccw+cw_, tan_para–ccw+cw_, tan_peri–ccw+cw_) where the x-axis represents the onset of the flashed probes from saccade onset, while the y-axis represents the sensitivity changes at a given retinotopic location. To then avoid any erroneous sensitivity results due to low sampling, these ten radial and tangential retinotopic functions were truncated from 540ms prior to saccade onset to 130ms after saccade onset, a temporal window which extends from (i.) pre-planning, (ii.) eye-movement planning, S_prep_, (computed by taking the temporal difference of S_onset_ from Go_onset_, central movement cue), and (iii.) execution (S_onset_ to S_offset_) phases of the cued detection task.

#### Normalized sensitivity functions

Despite measuring the CSF at the mean probe location for each experimental condition, it is possible that there are baseline sensitivity differences across probe locations within a condition. To therefore control for any eccentric-dependent effect on the raw sensitivity functions, I normalized these functions by the average sensitivity over the initial 50ms in the data (540ms to 490ms prior to saccade onset).

#### Periodicity of sensitivity functions

To investigate the frequency components of the sensitivity data, I first de-trended the data by subtracting the mean sensitivity (Supplementary Fig) **4B**. and then applied a fast Fourier transform (FFT) on the detrended data. To further investigate whether the observed periodicities included a spatial component (i.e., were spatially dependent), I randomly shuffled the probe location identities across experimental trials and calculated the sensitivity function and periodicity of the shuffled data using the same procedures as above. The shuffling procedure was repeated 1500 times for each experiment. Finally, I performed a set of two-tailed paired-sample t-tests between the FFTs calculated from the detrended data and the shuffled data to determine statistical significance.

## VI. ACKNOWLEDGMENTS

Thank you to Arnivan Nandy and Zixuan Xiao for helping with the initial experimental phase of this project. Finally, a special thanks to Michael Scudder for implementing the experimental design.

