## Supplementary material for "A fundamental law underlying predictive remapping": Supp Fig 1

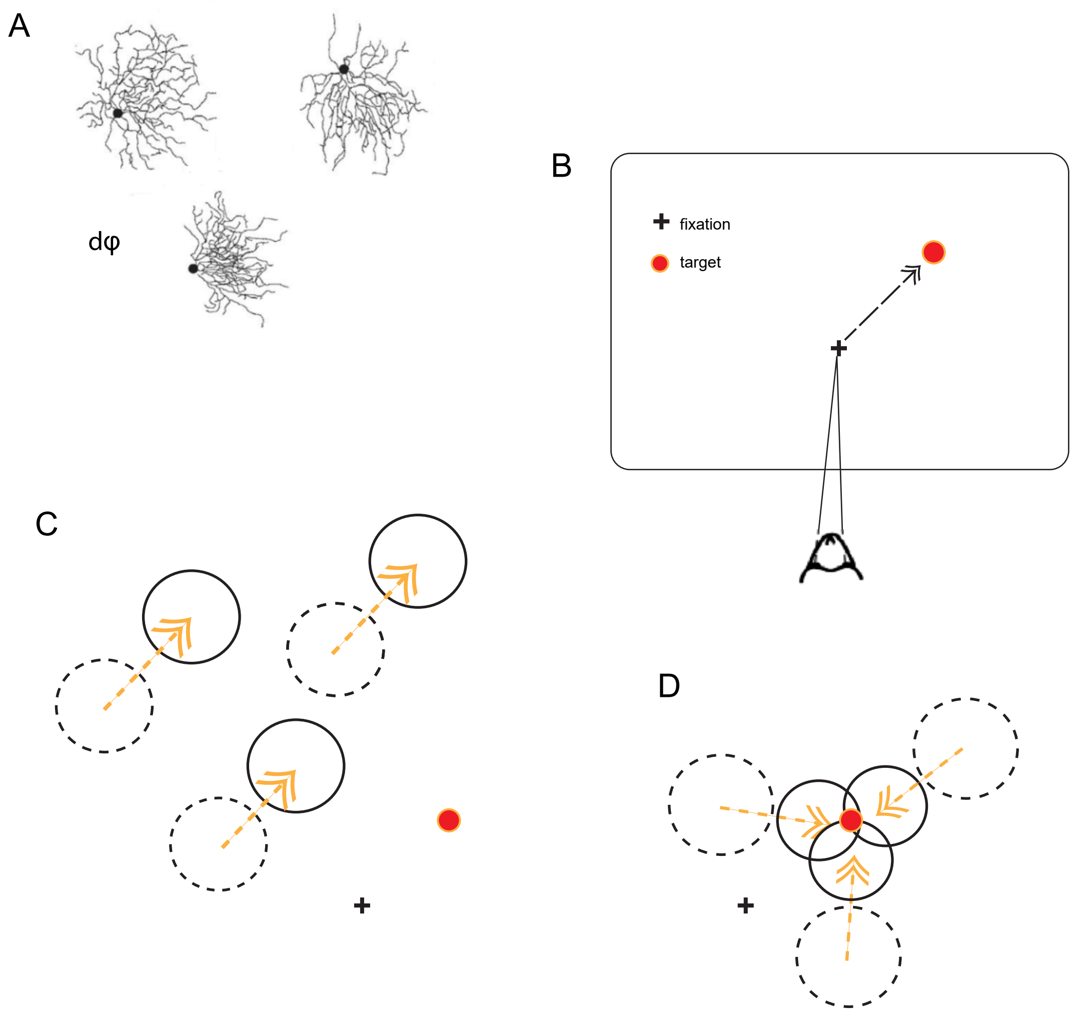


**Sup. Fig 1. Translational versus convergent form of predictive remapping**

(**A**) Cartoon of three cells in a dendritic field (dφ). A best-fit contour that includes the cell's center and dendritic extent is referred to the cell's anatomical classical receptive field (cRF) [see Ref.46]. (**B**) Cartoon of a foveated animal fixating and preparing to make a saccadic eye movement (black dotted arrow) towards an ecologically relevant visual target (red dot). (**C**) According to the translational form of remapping, cRFs (dotted black circles) do not respond when a pre-saccadic flashed probe (e.g., Gabor probe) is placed within these extents just before the saccade is made. Instead they respond when the flash occurs within their future post-saccadic locations (solid circles). This extent is roughly proportional the magnitude of the impending saccadic eye movement vector. (**D**) According to the convergent form of remapping the region around the target constitutes where neural responses will be the highest as if cRFs have converged around the pre-selected target.
