## Supplementary material for "A fundamental law underlying predictive remapping": Supp Fig 2

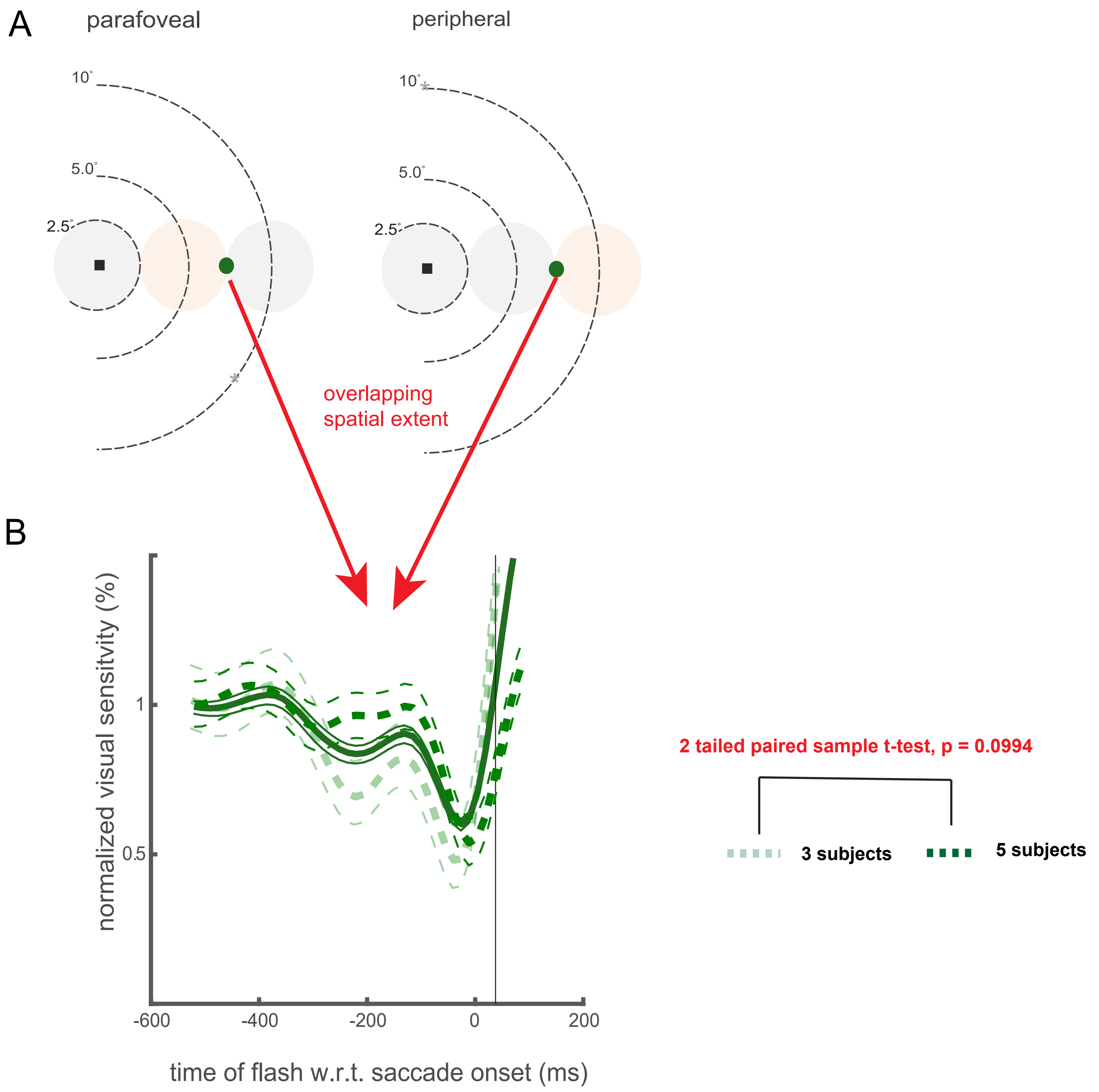


**Sup. Fig. 2. Sensitivity dynamics cannot be attributed  to different subject groups.**

(A) The overlapping point where sensitivity was measured for two different groups of subjects. (B) normalized changes in visual sensitivity as a function of flashed probe times relative to saccade onset between (dotted markers) and across (sold maker) the two groups. represent the error estimates calculated between groups.
