## Supplementary material for "A fundamental law underlying predictive remapping": Supp Fig 3

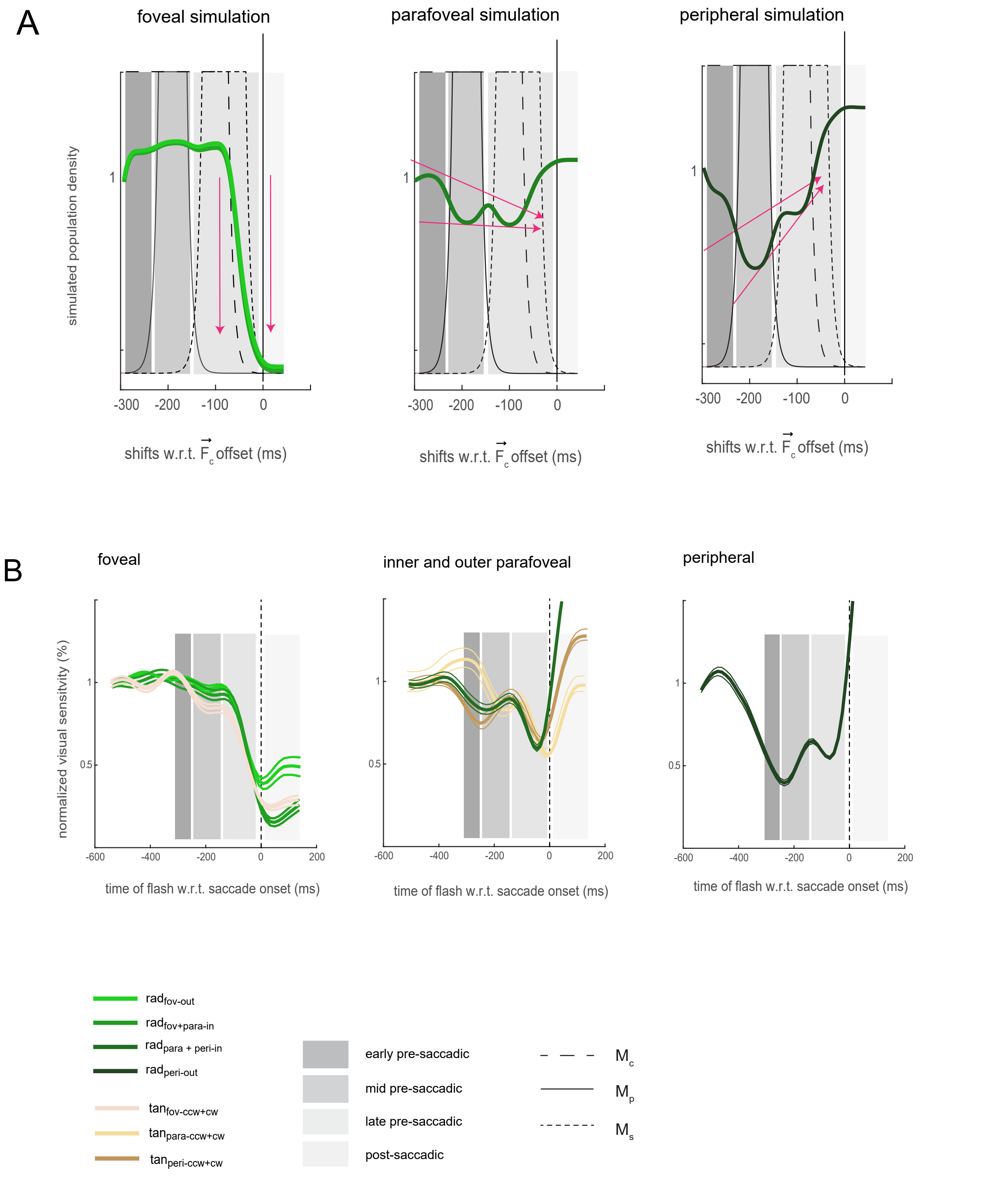


**Supplementary Fig. 3.** Sensitivity functions across visual space within distinct pre and post saccadic windows. A) Simulated population neural density. The arrows represent the general trend each sensitivity signature follows which mirrors the general signatures along radial and tangential axes observed in the behavioural experiments. B) Sensitivity functions across visual space from all three peripheral experiments.
