## Supplementary material for "A fundamental law underlying predictive remapping": Supp Fig 4

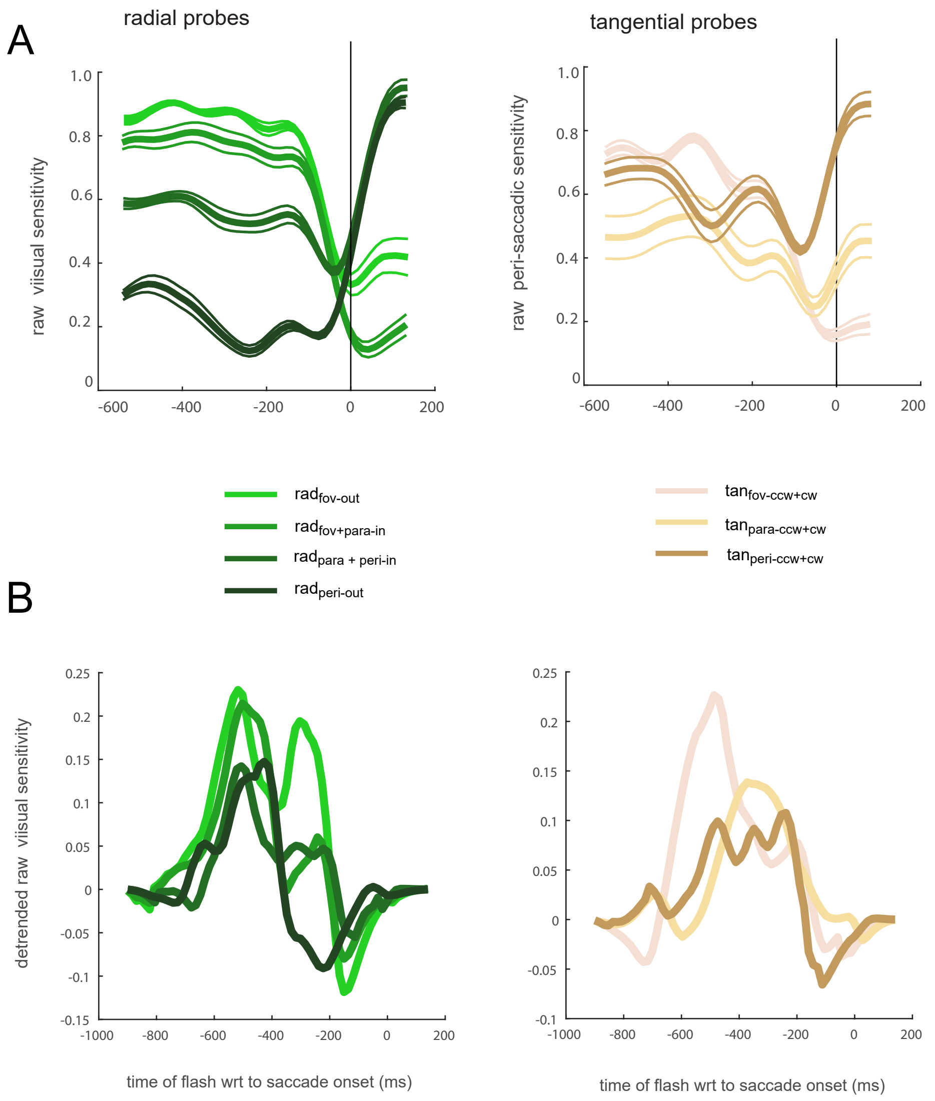


**Supplementary Fig. 4. Raw, detrended visual sensitivity**
(A) Raw sensitivity along the radial axis (left panel) and the tangential axes (right panel).
(B) Detrended sensitivity with the application of an Hanning window along the radial axis (left panel) and the tangential axes (right panel).
